# BacPROTACs mediate targeted protein degradation in bacteria

**DOI:** 10.1101/2021.06.09.447781

**Authors:** Francesca Ester Morreale, Stefan Kleine, Julia Leodolter, Stepan Ovchinnikov, Juliane Kley, Robert Kurzbauer, David M. Hoi, Anton Meinhart, Markus Hartl, David Haselbach, Markus Kaiser, Tim Clausen

## Abstract

Hijacking the cellular protein degradation system offers unique opportunities for drug discovery, as exemplified by proteolysis targeting chimeras (PROTACs). Despite their superior properties over classical inhibitors, it has so far not been possible to reprogram the bacterial degradation machinery to interfere with microbial infections. Here, we develop small-molecule degraders, so-called BacPROTACs, that bind to the substrate receptor of the ClpC:ClpP protease, priming neo-substrates for degradation. In addition to their targeting function, BacPROTACs activate ClpC, transforming the resting unfoldase into its functional state. The induced higher-order oligomer was visualized by cryo-EM analysis, providing a structural snapshot of activated ClpC unfolding a protein substrate. Finally, degradation assays performed in mycobacteria demonstrate *in vivo* activity of BacPROTACs, highlighting the potential of the technology to provide next generation antibiotics.

## INTRODUCTION

Technological advances in proteomics, chemical biology and high throughput screening campaigns have boosted drug discovery across therapeutic areas. Unfortunately, however, these advances have thus far not equally translated into the development of novel antibacterial agents (Payne et al., 2007; Tommasi et al., 2015). This discrepancy is most obvious when considering the small number of antibiotics that have been discovered in the last 50 years (Lewis, 2020). Beyond economic hurdles, the advancement of new antibiotics is challenged by the low permeability of the bacterial envelope, and the limited number of microbial proteins that can be specifically inhibited without off-target effects. The difficulties in finding effective anti-microbials are further compounded by the speed at which pathogens are developing resistance to existing drugs. In light of this unequal arms race, the return of bacterial pandemics is a real threat, and innovative strategies to combat infections by multi-resistant bacteria are urgently needed (Lewis, 2020).

An emerging concept in drug discovery is the induced elimination of target proteins. Engineered chemicals can now interfere with various degradation pathways, redirecting the lysosomal (Banik et al., 2020), the autophagy (Li et al., 2019; Takahashi et al., 2019) or the ubiquitin-proteasome systems (Sakamoto et al., 2001) to target specified proteins. The most prominent synthetic “degraders” are the Proteolysis Targeting Chimeras (PROTACs): bi-functional small molecules that contain a binding head for an E3 ubiquitin ligase and a chemical moiety to engage a protein of interest (POI) (Sakamoto et al., 2001). By bringing the E3 ligase and POI into proximity, PROTACs promote POI ubiquitination and consequent degradation by the proteasome. Protein degraders have various advantages over classical inhibitors. For example, they exhibit higher efficacy due to their catalytic mode of action. Moreover, they allow targeting of virtually any cellular protein and their modular architecture allows protein ligands to be repurposed to build degraders (Churcher, 2018; Hanzl and Winter, 2020; Schapira et al., 2019). Despite this promise, PROTAC technology is so far restricted to the ubiquitin tagging system of eukaryotes and has yet to be transferred to degradation pathways in bacteria. Fulfilling this last objective would be a powerful addition to the antibiotic arsenal.

Although ubiquitin is unique to eukaryotic cells, some bacteria utilize a similar system for targeted protein degradation. Phosphorylated arginine residues (pArg) serve as a degradation signal that is recognized by the ClpC:ClpP (ClpCP) protease, a quasi-proteasomal particle critical for microbial protein quality control, stress tolerance and pathogenicity. ClpCP is present in Gram-positive bacteria and in mycobacteria. In the latter, the equivalent ClpC1P1P2 protease is essential for survival *in vitro* and in macrophages (DeJesus et al., 2017; Rengarajan et al., 2005). ClpC carries out its quality control function by acting as an ATP-driven unfoldase that selects certain client proteins and translocates them into the protease compartment formed by ClpP. The ClpC protomer contains an amino-terminal domain (ClpC_NTD_), two AAA (ATPases Associated with diverse cellular Activities) domains termed D1 and D2, and a coiled-coil (M-domain) inserted into D1. In the active ClpC hexamer, the D1 and D2 domains form two stacked ATPase rings that power substrate unfolding and translocation. The ClpC_NTD_ on top of the D1 ring controls access to the unfoldase, providing docking sites for adaptor proteins including MecA (Schlothauer et al., 2003; Wang et al., 2011). In addition, the ClpC_NTD_ receptor domain recognizes pArg-tagged substrates that account for ~30% of the whole ClpP degradome in Gram-positive bacteria (Trentini et al., 2016). Compared to the eukaryotic proteasome, which recognizes a complex poly-ubiquitin signal, the degradation signal recognized by ClpCP is much simpler: a plain phosphate group attached to an arginine residue of the client protein (Trentini et al., 2016). Here we investigated whether the ClpCP degradation machine can be directly reprogrammed by small molecules. We reasoned that pArg-containing chemical adaptors could tether neo-substrates to the ClpC_NTD_ receptor domain, priming them for degradation (**Figure 1A**). Our results provide a proof of concept that such BacPROTACs can be developed and are active *in vivo*, enabling the inducible, selective and efficient degradation of target proteins in bacteria.

**Figure 1.**
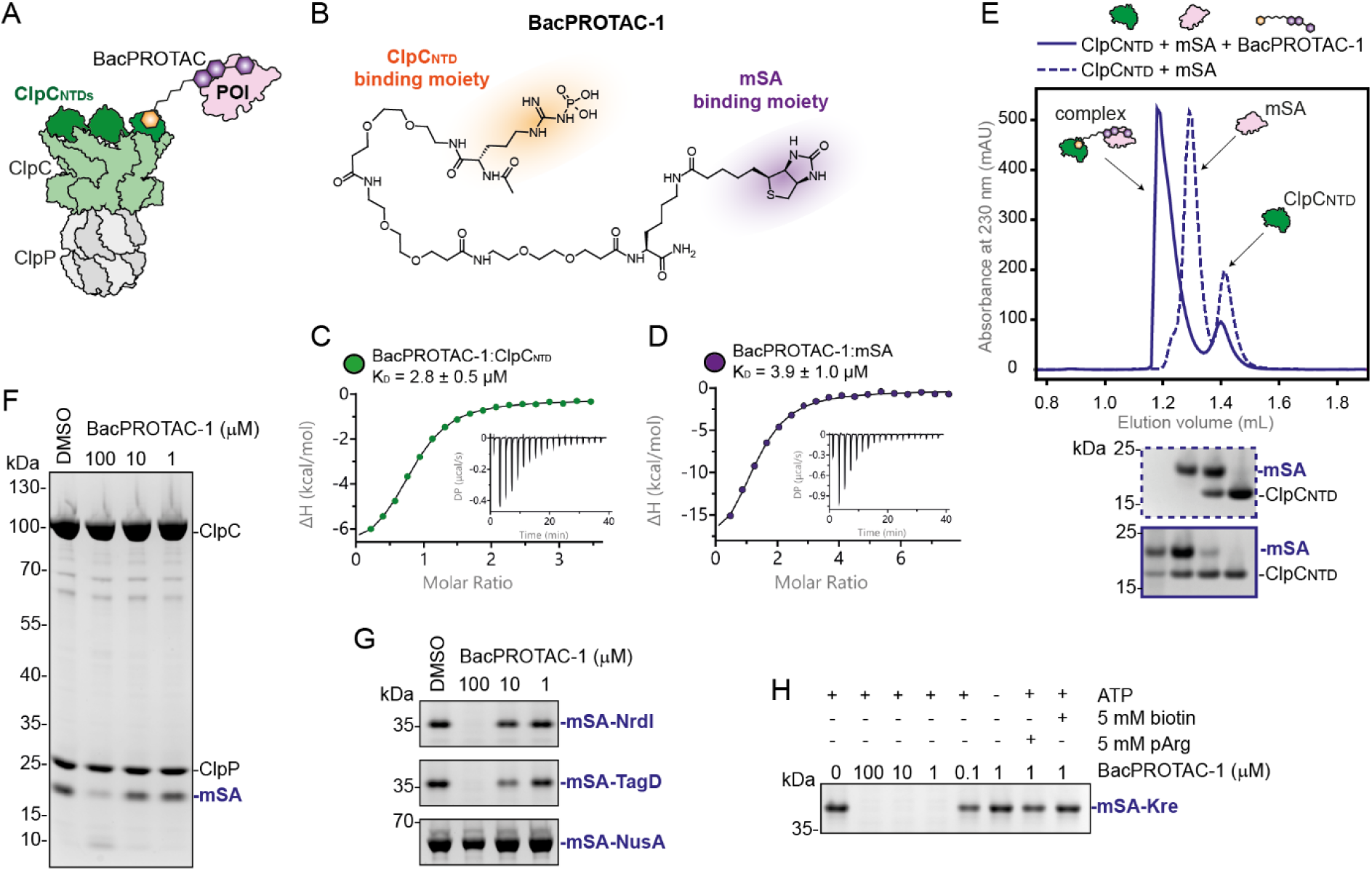
*In vitro* reprogramming of *B. subtilis* ClpCP by BacPROTAC-1. (**A**) Schematic representation of the BacPROTAC approach, hijacking the ClpCP protease. (**B**) Chemical structure of BacPROTAC-1, designed to tether the model substrate mSA to the ClpC_NTD_ receptor domain. (**C**, **D**) ITC titrations of BacPROTAC-1 against ClpC_NTD_ and mSA. Calculated K_D_’s and their standard deviation are indicated (*n* ≥ 3). (**E**) SEC analysis of a stoichiometric mSA:ClpC_NTD_ mixture in the presence (solid line) or absence (dashed line) of BacPROTAC-1. An SDS-PAGE analysis (lower panel) confirms the formation of the indicated ternary complex. (**F-H**) SDS-PAGE analysis of *in vitro* degradation assays. mSA alone and mSA fusion proteins (mSA-NrdI, mSA-TagD, mSA-NusA, mSA-Kre) are degraded by ClpCP in a BacPROTAC-1-dependent manner (2 hours incubation, DMSO used as control). (**H**) For mSA-Kre, further controls indicate that degradation is ATP-dependent and that pArg or biotin block the BacPROTAC-1-induced degradation. Uncropped images of SDS-PAGE gels are shown in **Figure S8**.

## RESULTS

### Development of BacPROTACs that reprogram ClpCP

To enable ClpC-mediated protein degradation, we designed BacPROTACs composed of a POI ligand, a chemical linker and a ClpC_NTD_ anchor. The anchor initially consisted of a peptidic pArg derivative mimicking the bacterial degradation tag. In order to test the targeting of a neo-substrate, we used monomeric streptavidin (mSA) as a model protein. We synthesized BacPROTAC-1 (**Figure 1B**), connecting the pArg moiety to biotin, a high affinity mSA ligand (Lim et al., 2013). The linker attachment points were designed based on high-resolution crystal structures of pArg:ClpC_NTD_ (Trentini et al., 2016) and biotin:mSA (Demonte et al., 2013). Isothermal titration calorimetry (ITC) measurements confirmed that BacPROTAC-1 binds mSA and ClpC_NTD_ with high affinity (K_DS_ of 3.9 μM and 2.8 μM **Figure 1C** and **1D**), whereas analytical size exclusion chromatography (SEC) runs revealed the formation of a stable ternary complex (**Figure 1E**). To analyze whether the induced spatial proximity is sufficient to trigger degradation, we reconstituted the *Bacillus subtilis* ClpCP protease *in vitro* and monitored mSA digestion at different BacPROTAC concentrations. Incubation with 100 μM BacPROTAC-1 led to selective mSA depletion (**Figure 1F**), indicating that the bacterial ClpCP can be reprogrammed by a pArg-containing chemical adaptor. Notably, however, efficient degradation was only observed at concentrations that were higher than the measured affinities among individual components. Presumably, this discrepancy reflects distinct requirements for substrate recruitment and translocation processes carried out by the ClpC unfoldase.

To follow up on this and analyze the influence of substrate-specific properties on ClpCP activity, we cloned various mSA fusion proteins. We selected four *B. subtilis* targets that have been identified as physiological, pArg-labelled ClpCP substrates (Trentini et al., 2016). Three of these proteins (NrdI, TagD, NusA) adopt compact protein folds, whereas Kre is predicted to be a mostly unstructured protein (**Figure S1**). *In vitro* assays, monitoring the BacPROTAC-1 dependent degradation of the four mSA fusion proteins by ClpCP, revealed pronounced differences (**Figure 1G** and **1H**). mSA-Kre was by far the best substrate, being degraded by 1 μM BacPROTAC. These *in vitro* data suggest that ClpCP preferentially acts on proteins with unstructured regions, which could serve as initiator sites for substrate unfolding and translocation. A similar mechanism has been proposed for the functionally related 26S proteasome, which requires an unstructured region to efficiently degrade ubiquitinated proteins (Inobe et al., 2011; Prakash et al., 2004).

To confirm that degradation is specifically induced by BacPROTAC-1, we hindered substrate recruitment by adding the isolated binding moieties pArg or biotin to the reaction (**Figure 1H**). In the presence of these compounds, mSA-Kre was not digested by ClpCP, confirming that BacPROTAC-mediated ternary complex formation is essential for degradation. Taken together, our results show that pArg-containing BacPROTACs can recruit POIs to the ClpC_NTD_ domain and promote their degradation by the ClpCP protease. In addition to the binding characteristics of the chemical adaptor, intrinsic properties of target proteins seem to play an equally important role in determining degradation efficiencies.

### BacPROTAC binding induces ClpC reassembly and activation

Housekeeping proteases and chaperones that target aberrant proteins need to be carefully controlled to prevent concomitant damage to functional proteins in the cell. A study addressing the regulation of the *Staphylococcus aureus* ClpC, a close relative to the *B. subtilis* unfoldase, revealed the existence of a ClpC resting state, a decamer with disrupted AAA rings (Carroni et al., 2017). Interaction with the adaptor protein MecA destabilizes the decamer and promotes assembly of the functional hexamer with an active arrangement of ATPase units (Carroni et al., 2017). In contrast to adaptor-mediated activation, it is unclear how the majority of ClpC substrates labelled with pArg trigger remodeling of the latent ClpC decamer. To elucidate this general activation mechanism – and more specifically, the way in which it is mimicked by a pArg-based degrader – we performed a structural analysis of ClpC in complex with BacPROTAC-1 and the mSA-Kre fusion protein. To stabilize ATP-mediated contacts between ClpC protomers, we used a catalytically inactive mutant (E280A/E618A, ClpC_DWB_) that binds ATP but does not hydrolyze it. When we added ClpC_DWB_ and mSA-Kre in stoichiometric amounts to a SEC column, ClpC_DWB_ and mSA-Kre eluted separately (**Figure 2A**). Like the *S. aureus* protein, isolated *B. subtilis* ClpC_DWB_ was present in its resting state, the decamer, as visualized by negative-staining EM (**Figure 2B** and **S2A**). Incubation with BacPROTAC-1 led to the co-elution of mSA and ClpC_DWB_, pointing to the formation of a stable ternary complex. To our surprise, however, the estimated size of the BacPROTAC induced complex was far from compatible with the predicted size of substrate-bound ClpC_DWB_ hexamer (**Figure 2A**). Instead the resulting complex behaved like a higher-order ClpC oligomer with a molecular mass beyond 2 MDa. Cryo-EM analysis of the substrate-engaged ClpC revealed the formation of a tetramer of ClpC hexamers, arranged in almost perfect tetrahedral symmetry (**Figure 2C** and **S2B**). A similar arrangement has been reported for a chimeric ClpC (*M. tuberculosis* NTD fused to *S. aureus* D1-D2) incubated with the antibiotic cyclomarin A (CymA) (Maurer et al., 2019). However, in that case, 2D class averages of negative-stained EM images could not reveal structural details of the tetrahedral assembly. BacPROTAC-tethered mSA-Kre, which mimics a trapped substrate, yielded seemingly better-defined particles of the activated state. Cryo-EM analysis visualized the overall organization of the higher-order ClpC unfoldase complex at 10 Å resolution (**Figure 2D** and **S2C, Table 1**). In this state, the D2 rings of the four ClpC hexamers project outwards such that they can interact with the ClpP protease. The substrate-bound ClpC_NTD_ domains are located in the center of the particle but are too flexible to be defined by EM density. Most strikingly, the four ClpC hexamers interact with each other via their coiled-coil M-domains, establishing a net of helix pairs holding the particle together (**Figure 2D**). As the M-domains are known to stabilize the resting state of ClpC (Carroni et al., 2017), their BacPROTAC-induced reorientation might disassemble the latent form and promote formation of active hexamers stabilized within a supramolecular assembly (**Figure 2E**). To resolve the functional units of the activated unfoldase at higher resolution, we performed a focused cryo-EM analysis of the single hexamers (**Figure 3A**). The resulting 3D map had an overall resolution of 3.7 Å (**Figure S2C-E** and **Table 1)** and allowed us to build an atomic model of the substrate-bound ClpC complex (**Figure 3**, **Movie S1**).

**Figure 2.**
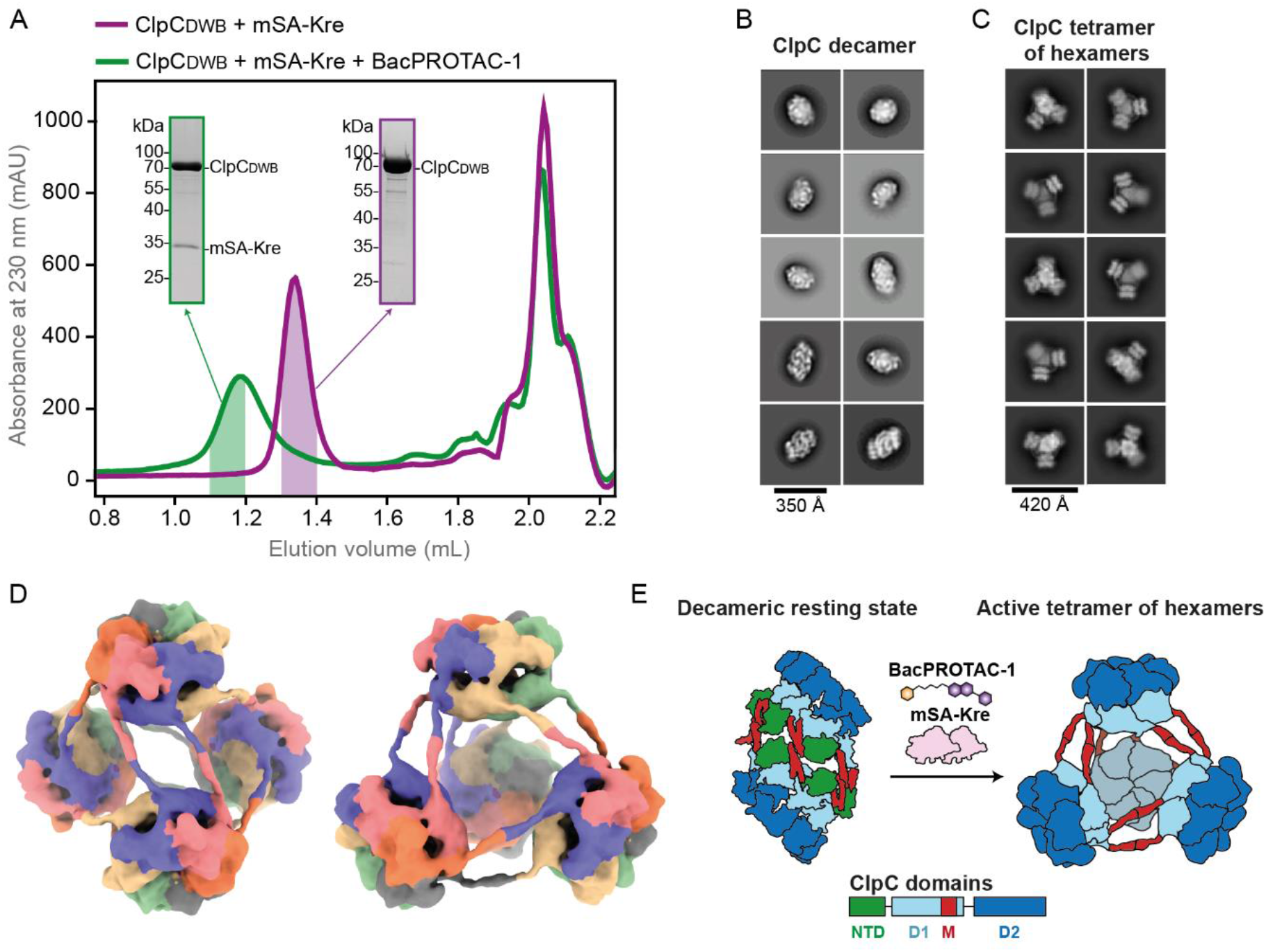
BacPROTAC-1 induces formation of an active ClpC oligomer. (**A**) SEC analysis of a stoichiometric ClpC_DWB_:mSA-Kre mixture in the presence (green) or absence (magenta) of BacPROTAC-1. The fractions used for EM analysis are highlighted. SDS-PAGE gels illustrate the coelution of ClpC_DWB_ and mSA-Kre in the presence of BacPROTAC-1. (**B**) Representative 2D class averages obtained from negative stained EM images, showing that *B. subtilis* ClpC forms a decameric complex representing the resting state. (**C**) Representative 2D class averages showing that in the presence of substrate and BacPROTAC-1, ClpC transforms into a 24-mer, composed of four hexamers present in functional form. (**D**) Refined 3D model (10 Å resolution) of the tetramer of ClpC hexamers, shown in two orientations. Individual ClpC protomers are colored differently. (**E**) Schematic representation of the BacPROTAC-induced conversion of the inactive ClpC decamer into the active higher-order particle (24-mer), using a domain-based coloring mode.

**Figure 3.**
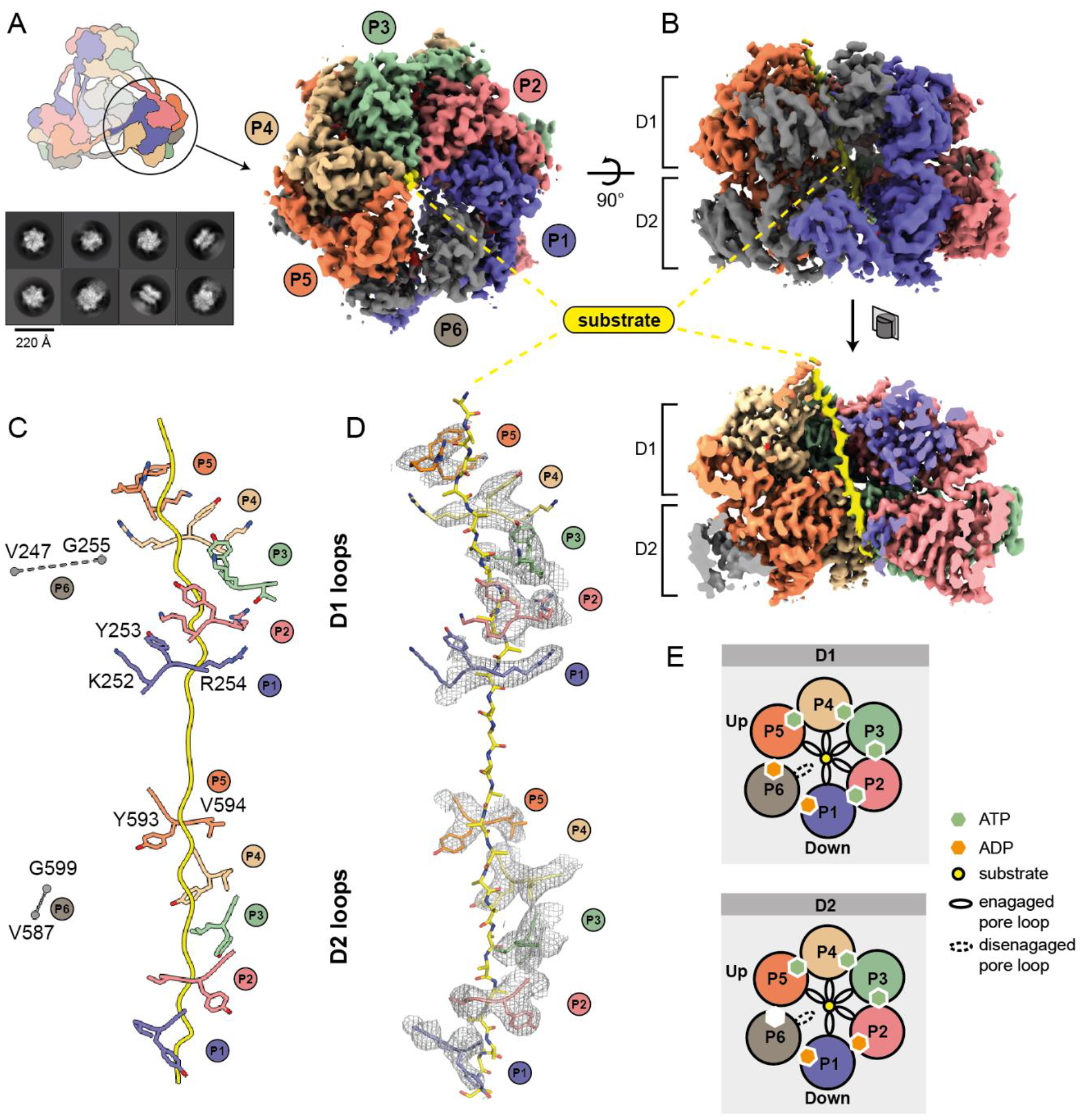
Cryo-EM structure of the activated ClpC hexamer in complex with a BacPROTAC-tethered substrate. (**A**) Representative 2D class averages are shown together with the final cryo-EM map, having a resolution of 3.7 Å. The density is colored according to subunits that are termed P1 to P6. The substrate captured in the central channel is shown in yellow. (**B**) Side view of the substrate-bound ClpC. The lower panel shows the cross section of the hexamer highlighting the substrate threaded through the two D1 and D2 rings of ClpC. The substrate was well defined by cryo-EM density over the entire passage of the central channel (80 Å). (**C**) Arrangement of primary D1 and D2 pore loops engaging the substrate (peptide backbone shown in yellow). The P6 pore loops, which were not in contact with the substrate, were too flexible to be modeled into the cryo-EM density. Their approximate position is indicated by flanking residues. (**D**) Cryo-EM density of the tyrosine bearing pore loops overlaid with the final model. (**E**) Schematic representation of nucleotide states and substrate engagement of the six ClpC protomers. Nucleotides were assigned based on cryo-EM density and distance matrices in the active site (**Figure S3**).

**Table 1.** Cryo-EM data collection, refinement and validation statistics.

|  | **ClpC hexamer**<br>**(EMDB-11707)**<br>**(PDB 7ABR)** | **Tetramer of CpC**<br>**hexamers**<br>**(EMDB-11708)** |
| --- | --- | --- |
| Data collection and processing |  |  |
| Magnification | 75,000 | 75,000 |
| Voltage (kV) | 300 | 300 |
| Electron exposure (e-/Å<sup>2</sup>) | 54 | 54 |
| Defocus range (µm) | 1.3-4.5 | 1.3-4.5 |
| Pixel size (Å) | 1.058 | 1.058 |
| Symmetry imposed | C1 | T |
| Initial particle images (no.) | 1,034,627 | 1,173,558 |
| Final particle images (no.) | 212,314 | 87,575 |
| Map resolution (Å) | 3.7 | 10 |
| FSC threshold | 0.143 | 0.143 |
| Map resolution range (Å) | 3.4 - 5.5 |  |
| Refinement |  |  |
| Initial model used (PDB code) | 3J3U |  |
| Model resolution (Å) | 3.7 |  |
| FSC threshold | 0.5 |  |
| Model resolution range (Å) |  |  |
| Map sharpening *B* factor (Å<sup>2</sup>) | -210 |  |
| Model composition |  |  |
| Non-hydrogen atoms | 27,209 |  |
| Protein residues | 3445 |  |
| Ligands | ADP: 4, ATP: 7 |  |
| *B* factors (Å<sup>2</sup>) |  |  |
| Protein | 67.93 |  |
| Ligand | 53.21 |  |
| R.m.s. deviations |  |  |
| Bond lengths (Å) | 0.011 |  |
| Bond angles (°) | 1.221 |  |
| Validation |  |  |
| MolProbity score | 2.33 |  |
| Clashscore | 19.31 |  |
| Poor rotamers (%) | 0.51 |  |
| Ramachandran plot |  |  |
| Favored (%) | 90.13 |  |
| Allowed (%) | 9.64 |  |
| Disallowed (%) | 0.24 |  |
| EMRinger score | 1.86 |  |

The most prominent feature of the cryo-EM reconstruction is a well-defined, 80 Å-long density that penetrates the entire ClpC pore, proceeding from the top of the D1 to the bottom of the D2 ring (**Figure 3B**). Although it is not possible to discern side chains, the density should represent the protein backbone of the captured mSA-Kre substrate, comprising 26 residues present in an extended conformation. The six ClpC subunits adopt a spiral arrangement, engaging the substrate in a similar manner to that observed in the related double-ring AAA unfoldases ClpA, ClpB, and Hsp104 (Gates et al., 2017; Lopez et al., 2020; Rizo et al., 2019). Five ClpC protomers (P1-P5, P1 as lowermost and P5 uppermost unit in the AAA ring) interact with the substrate through conserved tyrosine-bearing pore loops in the D1 and D2 domains, while the “seam” protomer (P6) is detached from the substrate, transitioning from the bottom to the top position of the spiral (**Figure 3C** and **3D**). Nucleotide states were assigned on the basis of the EM density and the position of the so-called arginine fingers from the neighboring subunit (**Figure S3**). Accordingly, the D1 and D2 ATPase rings act in a closely coordinated manner to translocate substrate through the central pore. In both rings, the substrate-engaging P2-P5 protomers are present in an ATP-bound state (except P2-D2), whereas protomers P1 and P6 accommodate ADP or are present in an apo state. Overall, the structural data are consistent with the previously suggested “hand-over-hand” translocation mechanism (Puchades et al., 2020), according to which nucleotide exchange is coupled with substrate release and upward movement of the seam protomer P6 to rebind ATP and substrate (**Figure 3E**). By reorienting in a concerted manner, the ClpC protomers move stepwise from one end of the spiral to the other, dragging the captured substrate down the central channel (Puchades et al., 2020). Cycles of substrate binding, translocation and release are driven by ATP hydrolysis at the lowermost P1 protomer of the spiral. In conclusion, the cryo-EM structure of the reconstituted ternary complex provides a snapshot of ClpC in the process of unfolding a BacPROTAC-tethered substrate. The structure of the ClpC 24-mer indicates that the pArg mark serves not only as a degradation signal, but also mediates higher-order oligomer formation and activation of ClpC. The developed BacPROTAC containing a pArg moiety triggers this remodeling mechanism and thus functions not only as a chemical adaptor, but also as an activator of the ClpCP protease.

### Extending the BacPROTAC approach to mycobacteria

The above biochemical and structural data revealed that pArg-based BacPROTACs can reprogram the ClpCP system of *B. subtilis*. To examine the therapeutic potential of the developed degrader, we aimed to transfer our approach to mycobacteria, which are among the most widely spread and dangerous human pathogens (World Health Organization, 2019). Although a pArg-dependent degradation pathway has not been identified in mycobacteria, the substrate receptor of the ClpC1P1P2 protease has a *bonafide* pArg-binding site, with all ClpC1_NTD_ residues implicated in binding the terminal phospho-guanidinium being conserved (**Figure S4**). To test if pArg-based BacPROTACs can reprogram the mycobacterial degradation machinery, we analyzed the *Mycobacterium smegmatis* Clp system. ITC and SEC experiments revealed that BacPROTAC-1 binds to ClpC1_NTD_ with high affinity (K_D_ = 0.69 μM) and promotes ternary complex formation with mSA and ClpC1_NTD_ (**Figure 4A** and **S5A**). We then reconstituted the ClpC1P1P2 protease *in vitro* and observed that BacPROTAC-1 induces the degradation of a mSA substrate in a highly selective and efficient manner (**Figure 4B**). These data demonstrate that pArg-containing degraders can reprogram the ClpC1P1P2 protease of mycobacteria.

**Figure 4.**
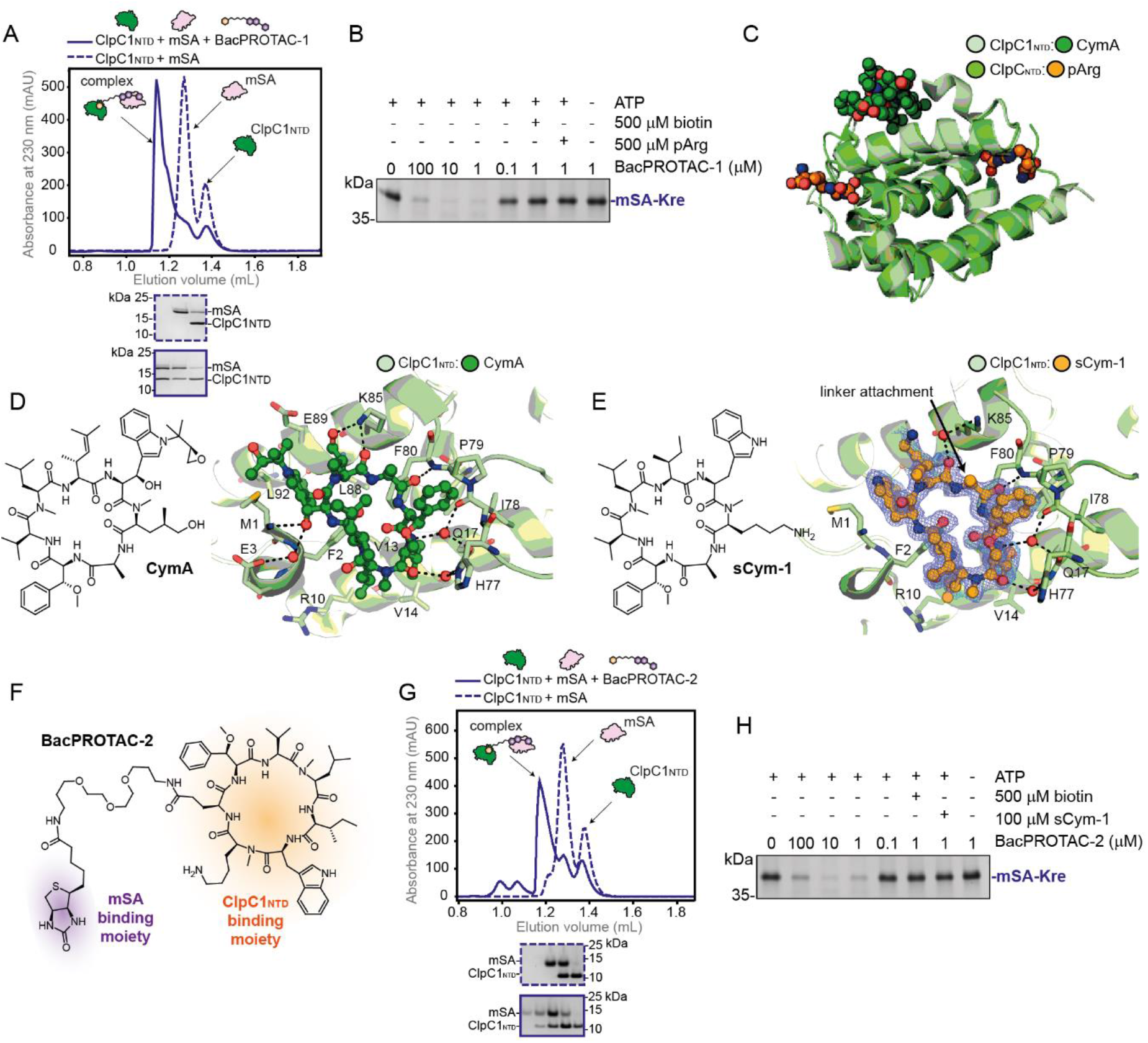
BacPROTACs can reprogram the mycobacterial ClpC1P1P2. (**A**) SEC analysis of a stoichiometric mSA:ClpC1_NTD_ mixture in the presence (solid line) or absence (dashed line) of BacPROTAC-1. SDS-PAGE analysis (lower panel) confirms the formation of the indicated ternary complex. (**B**) SDS-PAGE analysis of mSA-Kre degradation *in vitro*, following a 2-hour incubation with *M. smegmatis* ClpC1P1P2. The pArg-containing BacPROTAC-1 induces degradation in a concentration-dependent manner and can be outcompeted by separately provided pArg or biotin. (**C**) Superposition of ClpC_NTD_:pArg (PDB: 5HBN) with ClpC1_NTD_:CymA (PDB: 3WDC) crystal structures highlights the distinct locations of the ligand binding sites. (**D**) Chemical structure of CymA and co-crystal structure with ClpC1_NTD_ (PDB: 3WDC). Dotted lines represent hydrogen bonds, water molecules are shown as red spheres. (**E**) Chemical structure of the CymA analogue sCym-1 and co-crystal structure with ClpC1_NTD_ (PDB: 7AA4, this study), overlaid with the F_o_–F_c_ electron density map of the ligand (calculated at 1.7 Å resolution, contoured at 2σ). The arrow indicates the position for attaching the BacPROTAC linker. The respective alanine side chain does not contribute to the sCym-1:ClpC1_NTD_ interface. (**F**) Chemical structure of BacPROTAC-2, employing sCym-1 as a distinct ClpC1 anchor point. (**G**) SEC analysis of a stoichiometric mSA:ClpC1_NTD_ mixture in the presence (solid line) or absence (dashed line) of BacPROTAC-2. An SDS-PAGE analysis (lower panel) confirms the formation of the indicated ternary complex. (**H**) *In vitro* degradation of mSA-Kre in the presence of BacPROTAC-2, using the same assay conditions as in (A).

A major limitation in advancing pArg-based PROTACs is their poor pharmacokinetic profile and the chemical instability of the phospho-guanidinium group (Schmidt et al., 2014). To overcome these limitations, we looked for chemical entities that could replace pArg. Of note, the mycobacterial ClpC1 is targeted by a range of cyclic peptides that deregulate the ClpC1P1P2 complex (Lee and Suh, 2016). These antibiotics bind to the substrate receptor domain and thus have a pArg-like targeting function. The best characterized ClpC1_NTD_-directed antibiotic is CymA (Vasudevan et al., 2013), which is accommodated in a hydrophobic pocket located in a remote position relative to the pArg binding sites (**Figure 4C**). The CymA binding pocket is highly conserved in ClpC1 unfoldases, but is absent in ClpC proteins from Gram-positive bacteria, enabling selective targeting of the mycobacterial protease. Moreover, CymA can pass the cell envelope of mycobacteria (Schmitt et al., 2011) and thus represents a building block with important properties that could expand the potential of small molecule degraders in bacterial cells.

To prepare CymA-based degraders, we developed a solid-phase synthesis approach providing the 7-residue cyclic peptide in large quantities. To facilitate the intricate *de novo* synthesis of the natural compound (Barbie and Kazmaier, 2016), we replaced certain non-proteinogenic amino acids with chemically simpler analogues. Guided by structural data (**Figure 4D**) (Vasudevan et al., 2013), we prepared a series of CymA-like cyclic peptides and identified sCym-1 as a high affinity ClpC1_NTD_ ligand (K_D_ = 0.81 μM, **Figure S5B**). A co-crystal structure obtained at 1.7 Å resolution (**Figure 4E**, **Table S1**) confirmed that the sCym-1 mimetic adopts the same binding mode as the native CymA antibiotic. Moreover, the crystal structure revealed a possible linker attachment site to synthesize BacPROTAC-2, in which sCym-1 is linked to biotin (**Figure 4F**, **4G and S5C**). When tested in our *in vitro* assay, the sCym-1-based degrader stimulated removal of the mSA model substrate to similar extents as its pArg counterpart (**Figure 4H**), indicating that derivatives of the CymA antibiotic can be repurposed as BacPROTAC components. Importantly, these data also show that various ClpC1_NTD_ binders can be exploited to develop chemical adaptors for targeted protein degradation.

### BacPROTAC induces protein degradation in mycobacteria

We next analyzed whether BacPROTACs can induce POI degradation when introduced into mycobacteria. To show the system is generalizable, and since cellular biotin would compete for binding to mSA and impede BacPROTAC activity (**Figure 4H**), we looked for an alternative POI and identified the bromodomain-1 (BD1) of BRDT as attractive model substrate. BRDT_BD1_ (residues 21-137) encodes a soluble protein that binds with high affinity to BET bromodomain inhibitors (Matzuk et al., 2012). One of its small molecule ligands, JQ1, has been widely used in various PROTACs (Winter et al., 2015; Zengerle et al., 2015), thus facilitating the rational design of the bacterial degrader. Moreover, BRDT is a human protein with no bacterial homologues, allowing tracking of *in vivo* degradation without interfering with endogenous pathways. To target BRDT_BD1_, we synthesized BacPROTAC-3 linking sCym-1 and JQ1 (**Figure 5A**). The degrader was able to recruit BRDT_BD1_ (**Figure S5D-F**) and induce its degradation by ClpC1P1P2 in a highly specific manner (**Figure 5B** and **S6**). After validating the activity of the sCym-1 BacPROTAC-3 *in vitro*, we investigated whether it was able to reprogram ClpC1P1P2 in a cellular environment. For this purpose, we used *M. smegmatis* cells stably expressing BRDT_BD1_. We treated the culture with BacPROTAC-3 or alternatively, the individual building blocks sCym-1 and JQ1 (**Figure 5C**). After a 30-minute incubation, we quantified BRDT_BD1_ levels using capillary western blots. We found that BacPROTAC-3 induces BRDT_BD1_ degradation in a concentration-dependent manner, while sCym-1 or JQ1 treatments did not significantly alter BRDT_BD1_ levels (**Figure 5D** and **S7**). Given the reported deregulation of ClpC1P1P2 by CymA (Maurer et al., 2019), we next assessed whether BacPROTAC-3 led to selective elimination of BRDT_BD1_ or had a global impact on the mycobacterial proteome. To address this point, we performed a tandem mass tag mass spectrometric (TMT-MS) analysis of *M. smegmatis* lysates.

**Figure 5.**
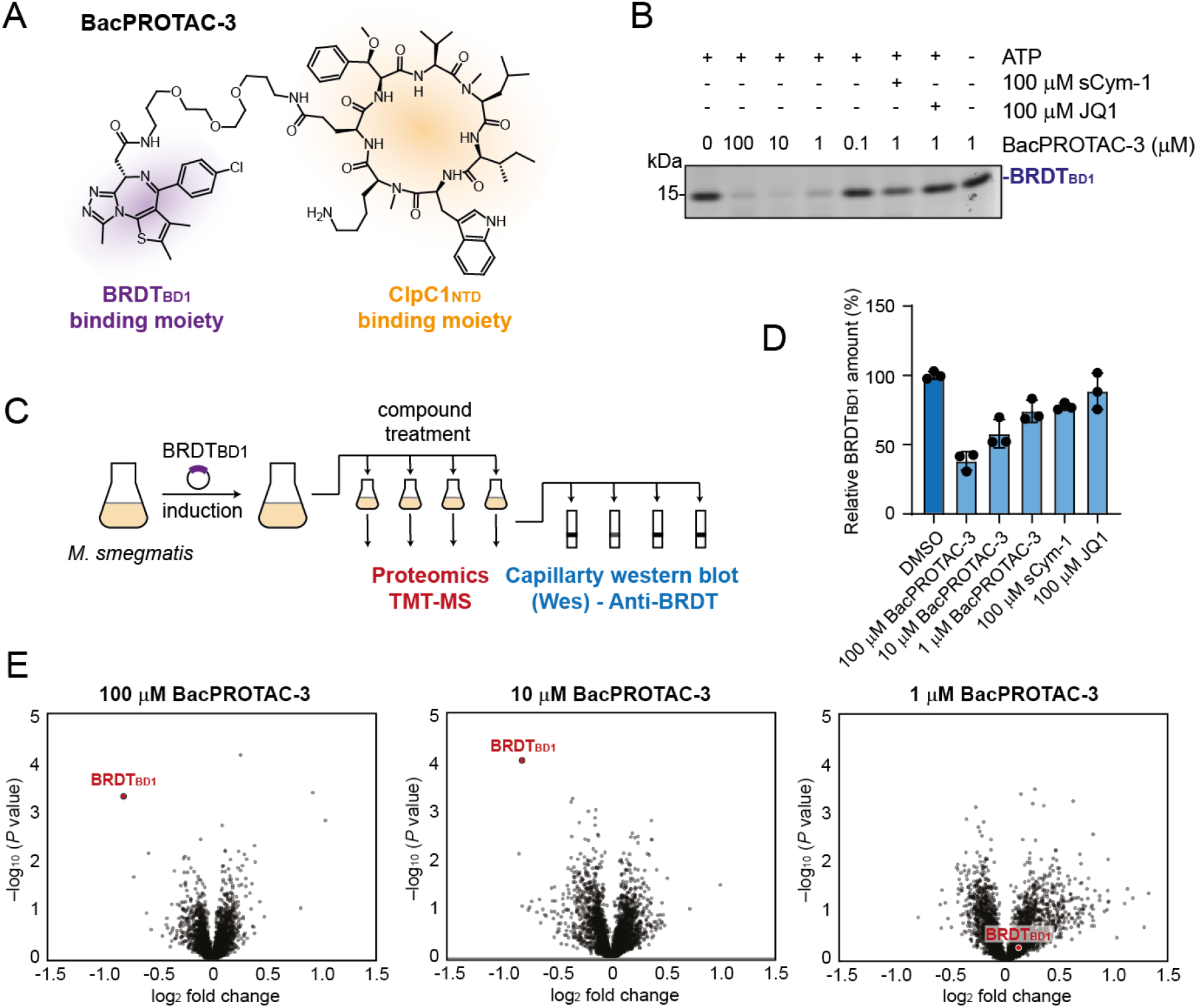
BacPROTAC-3 mediates degradation of a POI in mycobacteria. (**A**) Chemical structure of BacPROTAC-3 bridging JQ1 and sCym-1. (**B**) SDS-PAGE analysis of *in vitro* degradation following a 2-hour incubation with *M. smegmatis* ClpC1P1P2. BacPROTAC-3 promotes degradation of BRDT_BD1_ in a concentration-dependent manner and can be blocked by addition of its individual head groups. (**C**) Outline of BRDT_BD1_ degradation assay in mycobacteria. (**D**) Capillary Western blot (Wes) analysis of BacPROTAC-3 mediated effects on BRDT_BD1_ in the cell after 30 minutes of incubation. The bar chart summarizes three independent experiments, normalized to BRDT_BD1_ levels after DMSO treatment (dark blue) and plotted as mean ± SD. (**E**) TMT-MS proteomics analysis of the BacPROTAC-3 effect after 2 hours of incubation. The volcano plots show the fold-change (log2) in protein abundance in comparison to DMSO treatment, plotted against *P* value (–log_10_) (two-tailed Student’s T-test; triplicate analysis). The BRDT_BD1_ protein is highlighted in red.

Isobaric labelling allowed the detection and quantification of 2912 proteins. Among these, only BRDT_BD1_ was significantly (p < 0.001) depleted upon BacPROTAC-3 treatment (**Figure 5E**). Thus, the quantitative TMT-MS analysis provides compelling evidence that BacPROTAC-3 is capable of inducing degradation of the BRDT_BD1_ substrate in a highly specific manner. Together, our *in vivo* data demonstrate that CymA-based BacPROTACs can reprogram ClpC1P1P2 to induce POI degradation in mycobacteria. Furthermore, our data show that despite its chemical modification, the CymA scaffold maintains its ability to pass the mycobacterial cell envelope, providing an attractive tool for BacPROTAC development.

## DISCUSSION

Since resistance of pathogenic bacteria to current treatments is outpacing discovery of new antibiotics, it is crucial to expand the number of effective antimicrobial drugs. In the present work, we pioneered a methodology to develop next generation antibiotics. We developed small-molecule degraders that extend the PROTAC approach to bacteria. The resulting BacPROTACs reprogram the pArg-dependent ClpCP protease, the functional equivalent of the eukaryotic proteasome, and target it toward a specific substrate. As seen for pArg-and CymA-derived anchors, various molecules that bind to the substrate receptor of the ClpCP protease can be incorporated into BacPROTACs. By using a cell-permeable degrader, we also demonstrate that recruitment of a model protein to the mycobacterial ClpC1P1P2 protease leads to selective protein degradation in the bacterial cell. This *in vivo* analysis emphasizes the feasibility of applying the BacPROTAC approach to pathogens. The ClpC-directed degraders can be applied to Gram positive bacteria and mycobacteria, and can thus target pathogens that are a serious global health threat, including *M. tuberculosis* and Methicillin-resistant *S. aureus* (Lewis, 2020). Given the similar structural organization of the ClpAP protease, which also employs the N-terminal domain of the double ring ATPase ClpA as a substrate receptor, we presume that the BacPROTAC strategy presented here can be expanded to related Clp proteolytic complexes and used to combat the most severe pathogens across bacterial phyla.

Analogous to PROTACs active in eukaryotic cells, the bacterial degraders should have certain benefits over standard antibiotics that inhibit a limited group of microbial enzymes. The modular scaffold of BacPROTACs, for example, could allow many different POIs to be targeted for degradation. Currently, over 90,000 compounds directed against more than 70 different *M. tuberculosis* proteins are reported in the CHEMBL database. Connecting these POI ligands to ClpC1 binders would allow systematic evaluation of their potential to induce degradation of specific *M. tuberculosis* targets. The incorporated POI ligands do not need to occupy a functional site, and can thus target transcription factors, structural proteins and enzymes alike. High affinity binders that were identified in previous drug discovery efforts, but failed to block protein function, could therefore be repurposed as BacPROTAC building blocks and increase the number of druggable bacterial targets. Importantly, the developed approach could be employed to target virtually any bacterial protein since host-pathogen selectivity is achieved by reprogramming the ClpCP protease, which is absent in human cells. Beside targeting essential bacterial proteins, the developed approach could be used to eliminate certain virulence proteins. Reducing the evolutionary pressure to develop antibiotic resistance would be particularly interesting in combinatorial treatments with established antimycobacterial drugs. It should be also noted that the bacterial degraders seem to have a mechanistic advantage over their eukaryotic analogs. While PROTACs need to manipulate the complex E1-E2-E3 ubiquitination system to assemble a certain ubiquitin chain on a properly located lysine residue of the substrate, the bacterial degraders operate in a more straightforward manner. They influence the degradation reaction itself, bringing substrates directly to the ClpCP protease. We thus presume that optimizing degrader efficiency and finding suitable substrates could be facilitated, streamlining the rational design of potent antibacterial compounds.

The possibility of eliminating a POI from the cell by addition of a chemical compound has broad applications for basic research as well. The function of proteins in prokaryotic and eukaryotic cells is often investigated by studying knock-out strains. However, genetic manipulation is difficult when dealing with essential proteins, and requires special strategies that allow conditional, reversible and temporal control of protein levels. Chemical biology approaches that leverage the potency of small molecule degraders help to overcome these limitations (Burslem and Crews, 2020). Prominent examples are the auxin-inducible degron (AID) (Nishimura et al., 2009) and the selective removal of HaloTag fusion proteins (Neklesa et al., 2011). Likewise, the selective degradation of BRDT_BD1_-POI fusion proteins by BacPROTACs represents an attractive approach to generate conditional knockdowns in mycobacteria, allowing to dissect the function of essential proteins. These essential proteins may also be the very factors that are likely to become viable and effective targets for novel antimicrobials. In addition to addressing the biology of important microbial proteins, the degraders could thus help to validate antibiotic targets. As seen for small-molecule degraders in eukaryotic cells, we anticipate that BacPROTACs have the potential to accelerate drug discovery against pathogenic bacteria and facilitate the rational design of antibiotics targeting a wide range of essential microbial targets that were previously considered undruggable.

## Acknowledgements

We thank all members of the Clausen group for remarks on the manuscript and our colleagues Jan-Michael Peters and Yasin Dagdas for providing feedback on this study. Editing support was provided by Life Science Editors. **T**his work was supported by the European Research Council (AdG 694978, T.C.), an FFG Headquarter Grant (No 852936, T.C.), the Vienna Science and Technology Fund (WWTF LS17-029, F.E.M.) and Boehringer Ingelheim (Research Beyond Border program, J.L.). F.E.M. and J.L. are members of the Boehringer Ingelheim Discovery Research global post-doc program. The IMP is supported by Boehringer Ingelheim.

## Author contributions

F.E.M., A.M., M.K. and T.C. designed experiments; S.K. performed the chemical synthesis of the BacPROTACs, F.E.M. performed biochemical assays and binding measurements; F.E.M and D.H. performed the cryo-EM analysis, A.M. and J.K the crystallographic analysis; J.L., S.O. and R.K. prepared bacterial strains and carried out the *in vivo* analysis; D.M.H. and M.H. performed the mass spectrometry analysis; T.C. coordinated the research project and prepared the manuscript together with F.E.M.

## MATERIALS AND METHODS

### DNA constructs

Cloning of constructs for *E. coli* overexpression of full-length *B. subtilis* ClpC, ClpC_DWB_ (E280A/E618A), ClpC_NTD_ (1–148), ClpP, and BRDT_BD1_ was previously described (Filippakopoulos et al., 2012; Trentini et al., 2016). In case of expression constructs of *B. subtilis* ClpC, ClpC_DWB_, ClpC_NTD_ and ClpP, all variants were fusion constructs containing a C-terminal hexahistidine tag, except that for BRDT_BD1_ which contained a N-terminal hexahistidine tag followed by a TEV cleavage site.

Synthetic DNA of *M. smegmatis clpP1, clpP2* and *clpC1* genes (MSMEG_4673, MSMEG_4672, MSMEG_6091) was ordered from GeneArt (Thermo Fisher) and cloned into a pET21a vector DNA. ClpP1 and ClpP2 expression constructs encode a C-terminal tetrahistidine tag fusion, while that of ClpC1 expresses wild type protein. The coding region of *M. smegmatis clpC1* NTD (1–148) was cloned into pET21a and encodes a C-terminal hexahistidine tag fusion.

DNA for mSA-fusion constructs of three *B. subtilis* proteins (mSA-NrdI, mSA-TagD, mSA-NusA) cloned into a pETM14 vector was purchased from Genewiz. Expression constructs of mSA and mSA-Kre were cloned into a pNIC28-Bsa4 vector. These constructs encode a N-terminal hexahistidine tag followed by a TEV cleavage site fusion and a glycine-serine linker introduced between mSA and the bacterial protein. The mSA-Kre construct used for cryo-EM analysis was cloned into pET21a vector DNA and encodes a C-terminal hexahistidine tag fusion (**Table S2** reports the translated amino acid sequences of all mSA-fusion proteins).

Plasmid DNA for BRDT_BD1_ expression in *M. smegmatis* was ordered from Genescript. It contains the coding region of BRDT_BD1_ cloned into pMyC vector DNA. pMyC was a gift from Annabel Parret & Matthias Wilmanns (Addgene plasmid # 42192; http://n2t.net/addgene:42192; RRID:Addgene_42192).

### Protein expression and purification

Plasmid DNA was transformed into *E. coli* BL21 (DE3) or Rosetta cells (for *M. smegmatis* ClpC1, ClpP1 and ClpP2 proteins) and grown in LB broth supplemented with the respective antibiotic at 37 °C. Protein expression was induced by adding 0.1-0.5 mM isopropyl-1-thio-β-D-galactopyranoside (IPTG) at an OD_600_ of 0.8. Cells were cultured overnight at 18-20 °C, harvested by centrifugation and either lysed by sonication in a buffer containing 500 mM NaCl, 50 mM Tris pH 7.5, 10 mM imidazole, and 0.25 mM tris(carboxyethyl)phosphine (TCEP), or flash frozen and stored at −80 °C until purification.

Cell debris was removed by centrifugation. His-tagged proteins were purified from the cleared supernatants using Ni-NTA Agarose beads in batch. After several washing steps, bound protein was eluted using 50 mM Tris pH 7.5, 100 mM NaCl, 300 mM imidazole, 0.25 mM TCEP. Eluted protein was loaded onto a size exclusion chromatography column (Superdex 75 16/60 or Superdex 200 16/60 (GE Healthcare) depending on protein size) equilibrated in 50 mM Tris pH 7.5, 100 mM NaCl. For ClpC and ClpC_DWB_ purifications, the salt concentration of the size exclusion buffer was increased to 300 mM NaCl. Purified fractions were pooled and concentrated before flash freezing and stored at −80 °C.

Cell pellets for ClpP1 and ClpP2 purification were resuspended in 50 mM HEPES-NaOH pH 7.8, 300 mM NaCl, 30 mM imidazole and lysed by sonication. Cleared lysates were purified by Ni-NTA affinity chromatography (elution buffer: 50 mM HEPES-NaOH, pH 7.8, 300 mM NaCl, 250 mM imidazole) and subsequently by gel filtration using a Superose 6 16/60 column (GE Healthcare) equilibrated with 50 mM HEPES-NaOH pH 7.2, 150 mM KCl. 10% (*v/v*) glycerol was added to the elution fractions before flash freezing and storage at −80 °C. Processing of the full-length ClpP1 and ClpP2 to the mature ClpP1P2 complex was performed similar as previously described by Leodolter et al. (Leodolter et al., 2015).

Cell pellets for ClpC1 purification were resuspended in 50 mM Tris pH 7.5, 75 mM KCl, 2 mM EDTA, 10% (*v/v*) glycerol and lysed by sonication. After clarification of the lysate, ClpC1 was precipitated using 40% (*w/v*) ammonium sulphate. The pellet was resuspended in lysis buffer, loaded on a HiLoad 26/10 Q Sepharose High Performance column (GE Healthcare) equilibrated with lysis buffer and eluted in a gradient to 1 M KCl. ClpC1-containing fractions were pooled and precipitated again using 40% (*w/v*)ammonium sulphate. The pellet was resuspended in 50 mM HEPES-NaOH pH 7.2, 150 mM KCl, 10% (*v/v*) glycerol and loaded on a HiLoad 16/10 Superdex 200 prep grade gel filtration column (GE Healthcare) equilibrated in the same buffer. ClpC1 containing fractions were pooled and stored at −80 °C. Protein purity was monitored by Coomassie stained SDS-PAGE and correct molecular mass of purified proteins was verified by mass spectrometry.

### Isothermal Titration Calorimetry (ITC)

ITC experiments were performed using a MicroCal PEAQ-ITC instrument (Malvern) at 25 °C in a buffer containing 50 mM Tris pH 7.5, 100 mM NaCl. Each titration consisted of 19 injections with intervals of 120 s (the first injection of 0.4 μL was followed by 18 injections of 2 μL) at constant stirring at 750 rpm. DMSO concentration was matched between cell and syringe to be 2% (*v/v*). Data were fitted using a single binding site model with a fitted offset subtraction using the MicroCal PEAQ-ITC Analysis Software. Each titration was repeated at least twice.

Titrations involving BacPROTAC-1 were performed with the ligand loaded into the syringe and the protein into the cell. 400 μM BacPROTAC-1 was titrated against 20 μM ClpC_NTD_ or mSA. A control titration of ligand into buffer was performed in order to determine the heat of dilution.

Titrations involving sCym-1, BacPROTAC-2 and BacPROTAC-3 were performed with the ligand loaded into the cell and protein into the syringe to account for the low solubility of the compounds in aqueous solutions.

### Analytical size exclusion chromatography (SEC)

For the analytical runs involving ClpC/C1_NTD_ and mSA or BRDT_BD1_, the proteins were premixed at equimolar concentrations (25 μM) and BacPROTAC-1, BacPROTAC-2, BacPROTAC-3 (25 μM) dissolved in 100% (*v/v*) DMSO giving rise to a final concentration of 0.25% (*v/v*). For control experiments only DMSO was added to that concentration. Samples were loaded into a 30 μL loop and applied to a Superdex 75 3.2/300 increase column (GE healthcare) equilibrated in 50 mM Tris pH 7.5, 100 mM NaCl. Runs were performed at room temperature at a flow rate of 0.06 mL/min. Each run was performed in triplicate. 100 μL fractions were collected and analyzed by SDS-PAGE and Coomassie staining.

For electron microscopy analysis, *B. subtilis* ClpC_DWB_ (25 μM monomer) and the substrate mSA-Kre (25 μM) were premixed in a buffer containing 50 mM Tris pH 7.5, 50 mM KCl, 5 mM MgCl_2_, 5 mM ATP and 1.25 % (*v/v*) DMSO or 156 μM BacPROTAC-1 (thereby adding 1.25% (*v/v*) DMSO as vehicle). 30 μL were loaded on a Superose 6 3.2/300 increase column (GE Healthcare) equilibrated in 50 mM Tris pH 7.5, 50 mM KCl, 5 mM MgCl_2_, and 2.5 mM ATP. The experiment was performed at room temperature at a flow rate of 0.06 mL/min. 100 μL fractions were collected and used to prepare grids for EM analysis (**Figure 2A**).

### *In vitro* degradation assays

*In vitro* degradation assays containing 0.5 μM *B. subtilis* ClpC (hexameric), 0.5 μM *B. subtilis* ClpP (heptameric), 2 μM substrate protein, 15 mM phosphoenolpyruvate (PEP), 10 U/mL pyruvate kinase (Sigma Aldrich) were performed in a buffer containing 50 mM Tris pH 7.5 (at 37°C), 50 mM KCl, 20 mM MgCl_2_, 10% (*v/v*) glycerol. Compounds were dissolved in 100% (*v/v*) DMSO and further diluted giving rise to a final concentration of 1% (*v/v*) of DMSO in the assay. In control experiments DMSO was added to that concentration. Reactions were started by addition of 5 mM ATP and terminated by adding SDS sample buffer after 2 hours incubation at 37 °C. The samples were analyzed by SDS-PAGE and Coomassie staining. Degradation assays using 0.5 μM *M. smegmatis* ClpC1 (hexameric), 0.25 μM *M. smegmatis* mature ClpP1P2 (both heptameric), 2 μM substrate protein, 15 mM PEP, 10 U/mL pyruvate kinase (Sigma Aldrich) were carried out in 50 mM HEPES-NaOH pH 7.2, 100 mM KCl, 10 mM MgCl_2_, 10% (*v/v*) glycerol using the same procedure as described above. All experiments were performed in triplicates which gave similar results.

### Negative staining EM sample preparation, data collection and processing

In the absence of BacPROTAC-1, ClpC_DWB_ and the substrate mSA-Kre eluted separately in analytical SEC. The fraction containing ClpC_DWB_ (**Figure 2A**) was applied onto glow-discharged carbon-coated Cu/Pd Hexagonal 400 mesh grids (Agar Scientific) subsequently stained with a solution of 2% (*w/v*) uranyl acetate. The grids were screened and then imaged using a FEI Tecnai G2 20 microscope equipped with a 4k Eagle camera (FEI) using a pixel size of 1.8 Å/px. 1,004 micrographs were collected (a representative micrograph is shown in **Figure S2A**) and 856 particles were manually picked and 2D classified, generating templates for subsequent automatic particle picking on the entire set of micrographs. 369,415 picked particles were extracted using box size of 300 px and subjected to 2D classification. The resulting highest populated 2D class averages are shown in **Figure 2B**.

### Cryo-EM sample preparation and data collection

The ClpC_DWB_:BacPROTAC-1:mSA-Kre complex was isolated using analytical SEC. Cryo-EM grids were prepared using glow discharged R2/2 Cu 200 mesh grids (Quantifoil) pre-floated onto custom-made 2.9 nm continuous carbon film. 4 μL sample were applied onto a grid held in the sample grid chamber of a Leica EM GP instrument (Leica Microsystem Inc.) cooled at 4 °C, with a relative humidity of 80-90%. Grids were blotted 2 s with a Whatman type 1 filter paper using the blotting sensor and flash frozen in liquid ethane. The quality of the grids was screened using a Thermo Scientific Glacios Cryo Transmission Electron Microscope equipped with a Falcon3 Direct Electron Detector. For the best grid, a dataset of 4,455 micrographs was collected (a representative micrograph is shown in **Figure S2B**) on a Titan Krios equipped with a Falcon 3EC detector with a nominal magnification of 75,000 resulting in pixel size of 1.058 Å/px. A total dose of 54 e^-^/Å^2^ was fractionated into 40 frames. 452 poor quality images (heavy contamination, poor ice quality or bad information transfer) were discarded after visual inspection, leaving 4,003 images for further analysis.

### Cryo-EM analysis of the ternary complex between ClpC, BacPROTAC-1 and mSA: the tetramer of hexamers

Multi-frame micrographs were analyzed in cryoSPARC v2 (Punjani et al., 2017). The full micrographs were motion corrected and their contrast transfer function (CTF) parameters were estimated. 482 particles were manually picked and 2D classified, generating a template for subsequent automatic particle picking on the entire set of micrographs. 1,173,558 picked particles were extracted using box size of 500 px and subjected to two rounds of 2D classification, selecting only the classes containing the full tetramer of hexamers for further analysis (**Figure 2C**). 87,575 “clean” particles were used for *ab initio* model generation and subsequent homogeneous refinement applying T symmetry. The approximate resolution of the obtained map was 10 Å as judged by Gold Standard Fourier shell correlation (FSC) (**Figure S2C**).

### Cryo-EM analysis of the ternary complex between ClpC, BacPROTAC-1 and mSA: the hexameric building block

For the analysis of the single hexamers, the same multi-frame micrographs were motion corrected and dose-weighted using MotionCor2 1.0.5 (Zheng et al., 2017). Aligned micrographs were used for CTF estimation in Gctf 1.06 (Zhang, 2016). A subset of 10 micrographs was used for manual particle picking in crYOLO v1.3.5 (Wagner et al., 2019), the picking centers were the single hexamers rather than the tetramer of hexamers. Manually picked particles were used to train a model for automatic particle picking on the full micrographs set (160 px box size). A total of 1,034,627 auto-picked particles were imported in RELION 3.0 (Zivanov et al., 2018). Particles were extracted with 2 × 2 binning (box size 300 px, rescaled to 150 px) and subjected to two rounds of 2D classification. The particles belonging to the best 2D classes (**Figure 3A**) were re-extracted un-binned (box size 300 px) and 3D classified using an initial 3D model generated with cryoSPARC v2 (Punjani et al., 2017). After 3D classification, 212,314 particles were selected for 3D refinement. The resolution was estimated with Gold Standard FSC, and B-factor sharpening was applied, both using the RELION postprocessing. The final map used for model building had an overall resolution of 3.7 Å. Local resolution was estimated in RELION and is shown in **Figure S2E.**

### Model building

An initial model was built in Coot (Emsley et al., 2010) by rigid body fitting of secondary structure elements of one of the previously reported ClpC structures into the EM-map (PDB: 3J3U) (Liu et al., 2013). This model, however, required partial rebuilding and several steps of manual real-space refinement in Coot. Substrate-bound ClpB cryo-EM structures (Deville et al., 2019) were used as reference, as mSA-Kre binding caused substantial conformational rearrangements in the unfoldase hexamer when compared to the apo form of ClpC reported previously (Liu et al., 2013; Wang et al., 2011). The substrate polypeptide chain was modelled as a poly-alanine. Phenix (Afonine et al., 2018) was used for further real-space refinement. The quality of the model and model to map fitting were assessed using Phenix (Afonine et al., 2018), MolProbity (Williams et al., 2018) and EMRinger (Barad et al., 2015). Structural illustrations were prepared using either UCSF Chimera (Goddard et al., 2007), UCSF ChimeraX (Goddard et al., 2018), or Pymol (Schrodinger, 2015).

### ClpC1_NTD_:sCym-1 co-crystallization and structure determination

ClpC1_NTD_ in a buffer containing 10 mM Tris-Cl pH 7.5, 100 mM NaCl, 1 mM sCym-1, and 1% (*v/v*) DMSO was crystallized in a hanging drop, vapor diffusion setup at 15 mg/mL concentration using a reservoir solution containing 100 mM MES/imidazole pH 6.5, 6% (*w/v*) PEG 20K, 12% (*v/v*) PEG MME 550, 120 mM 1,6-hexanediol, 120 mM 1-butanol, 120 mM (*RS*)-1,2-propanediol, 120 mM 2-propanol and 120 mM 1,4-butanediol. Crystals were grown at room temperature for one week, subsequently harvested and flash-cooled in liquid nitrogen. Diffraction data were collected at an in-house X-ray generator and processed, scaled using the XDS (Kabsch, 2010) package to a resolution of 1.7 Å. Initial phases were obtained by Molecular Replacement using PHASER and the structure of ClpC1_NTD_ (PDB: 3WDB) (Vasudevan et al., 2013) as starting model. The model was improved in iterative cycles of manual building using Coot (Emsley and Cowtan, 2004) and refinement with Phenix (Liebschner et al., 2019) omitting 5% of randomly selected reflections for calculation of Rfree. Model quality was monitored using MolProbity (Williams et al., 2018) and the final model exhibited good stereochemistry with 97.5% of residues in favored regions of the Ramachandran plot and without any outliers. Structural illustrations were made using Pymol (Schrodinger, 2015).

### *In vivo* degradation assays

A stationary culture of *M. smegmatis* mc^2^-155 carrying a plasmid for BRDT_BD1_ expression was diluted 1:200 in 50 mL 7H9 medium supplemented with 50 μg/mL hygromycin. The culture was grown at 37°C under vigorous shaking to an OD_600_ of ~1. Expression of BRDT_BD1_ was subsequently induced by addition of 200 μL 50% (*w/v*) acetamide in H_2_O. The culture was maintained for additional 7 hours thereby reaching an OD_600_ of ~4. Cells were harvested by centrifugation and resuspended to an OD_600_ of 5 in fresh 7H9 medium. 270 μL aliquots of the culture were transferred in a 96-well glass-coated microplate (Thermo Scientific) and supplemented with either 1% (*v/v*) DMSO, 100 μM, 10 μM, or 1 μM BacPROTAC-3, 100 μM sCym-1, or 100 μM JQ1 (Note, the medium of contained final 1% (*v/v*) DMSO). Experiments were performed in triplicates. Aliquots of 250 μL of each cell suspension were withdrawn after 30 minutes incubation, cells were harvested, and the pellets were flash-frozen in liquid nitrogen. Cell pellets were thawed, resuspended in 100 μL resuspension buffer (50 mM HEPES-NaOH pH 7.2, 150 mM KCl) and lysed for 10 minutes using the Bioruptor (Diagenode, 10 cycles, 30 seconds on - 30 seconds off) in presence of small amounts of glass beads. Lysates were clarified by centrifugation, flash-frozen and stored at −80 °C.

Cytosolic amounts of BRDT_BD1_ and RpoB in the lysates were quantified using Wes (ProteinSimple). Bacterial lysates were diluted 2-fold and analyzed using the Protein Simple 12-230 kDa Wes Separation Module following the manufacturer’s instructions. Anti-BRDT (Bio-Vision) and anti-RpoB (BioLegend) antibodies were combined in a single primary antibody mixture for simultaneous BRDT_BD1_ and RpoB detection. The primary antibody mixture contained anti-BRDT diluted 1:250 and anti-RpoB diluted 1:25,000 in Protein Simple Wes Antibody Diluent 2. BRDT_BD1_ and RpoB were detected using the Anti-Mouse (Protein Simple) and Anti-Rabbit (ProteinSimple) detection modules for Wes following the manufacturer’s instruction. Anti-Mouse and anti-Rabbit antibodies were combined 1:9 and the obtained secondary antibody mixtures were used for detection. Results were analyzed using the Compass for SW software (ProteinSimple). Compass displays the chemiluminescent signal detected along the length of the capillaries as electropherograms, where the intensity of the chemiluminescent signal is plotted against the apparent molecular weight (**Figure S7**). Detected peaks were quantified and include a peak at the expected BRDT_BD1_ molecular weight (BRDT_BD1_ MW = 16.6 kDa) and an additional peak at the expected RpoB molecular weight (MW = 128.5 kDa). Peak areas were normalized to DMSO-treated cells and plotted as mean ± SD (**Figure 5D** and **S7G**).

### Sample preparation for quantitative mass spectrometry analysis

For quantitative MS analysis, *M. smegmatis* cells expressing BRDT_BD1_ were treated for 2 hours with 1% (*v/v*) DMSO, 100 μM, 10 μM, or 1 μM BacPROTAC-3, following the procedures described above. Cleared lysates were processed according to the single-pot SP3 protocol (Hughes et al., 2019) for low input proteomics sample preparation. Each lysate of 100 μl was reduced with 10 mM dithiothreitol (DTT, Sigma Aldrich) for 45 minutes at 37 °C and subsequently alkylated with 20 mM iodoacetamide (IAA, Sigma Aldrich) at room temperature for 1 hour. In parallel, a 1:1 mixture of 50 mg/mL Sera-Mag SpeedBeads (GE Healthcare) and 50 mg/mL Sera-Mag SpeedBeads (GE Healthcare), exhibiting different surface hydrophilicity, was prepared in water. To each lysate 15 μL of the prepared SP3 bead stock was added and binding was induced by the addition of 100 μL ethanol. To ensure proper binding, samples were incubated on a shaker for 5 minutes at 24 °C and 1000 rpm. After protein binding, beads were washed 3 times with 200 μL rinsing solution (80% (*v/v*) ethanol in water) while being kept on a magnetic rack. Protein elution from the beads was enforced by addition of 100 mM triethylammonium bicarbonate (pH = 8.5, Sigma Aldrich). To disaggregate the beads, the tubes were shortly sonicated in a water bath. For protein digestion 1:25 wt/wt ratio of trypsin to protein was added and the samples were incubated overnight at 37 °C in a thermo-shaker at 1,000 rpm.

Peptides were labelled for quantification in a multiplexed setup with TMT isobaric mass tags (TMTpro™ 16plex). Sample amount and quality was determined by HPLC-UV using a Dionex UltiMate 3000 HPLC RSLC nanosystem with a PepSwift Monolithic RSLC column (0.2 x 5 mm, Thermo Fisher Scientific) at 60 °C. Peptides were separated using a 20 minutes 2-90% elution gradient of buffer A (80% (*v/v*) ACN, 0.1% (v/v) TFA in aqueous solution). For the labelling procedure, one TMTpro set was split into 3 aliquots to label 3 replicates. 14 channels (TMTpro 126-133N Da) were distributed over 2 timepoints for each of the 7 treatments. The 2 remaining channels (133C and 134N) were used as reference channels with pools of all samples. Each sample was tagged with an excess of the respective TMT labelling reagent (1:20, peptide:TMT label) and incubated at room temperature for 1 hour. The reaction was quenched by addition of 5 μL 5% (v/v) hydroxylamine (Sigma Aldrich), followed by a 15 minutes incubation step. For each replicate all 16 channels were pooled, and the volume was reduced to 100 μL in a speedvac. Removal of excess TMT labelling reagent was achieved by running the samples through tips filled with silica gel equilibrated in water.

### High pH chromatography and LC-MS/MS analysis

Peptides were separated using a 40 min 2-50% gradient of buffer A in a high pH chromatography (TEA, pH = 8.5) setup using a Dionex UltiMate 3000 HPLC RSLC nanosystem with a XBridge Peptide BEH C18 Column (1 x 150 mm, 130 Å, 3.5 μm, Waters). 40 fractions were collected and pooled by combining every 11^th^ fraction to generate a final number of 10 fractions for each replicate. The volume of each sample was adjusted to 100 μL and the sample amount was estimated by running monolithic control runs.

LC-MS/MS analysis was performed on a Dionex UltiMate 3000 HPLC RSLC nanosystem using an Acclaim PepMap C-18 precolumn (0.3 x 5 mm, Thermo Fisher Scientific) and an Acclaim PepMap C-18 column (50 cm x 75 μm, Thermo Fisher Scientific) coupled to a Q Exactive HF-X hybrid quadrupole-Orbitrap mass spectrometer (Thermo Fisher Scientific). Peptides were separated using a 120 min linear gradient of 2-35% buffer A at a flowrate of 230 nL/min. MS1 spectra were generated in a 380-1,650 *m/z* mass range at a 120,000 orbitrap resolution, AGC target of 3e6, with a maximum injection time of 50 ms. The top 10 precursors were selected for MS2 analysis using a 0.7 *m/z* quadrupole precursor isolation window, allowing charge states 2-4 and a dynamic precursor exclusion of 30 s. The orbitrap was operated at 45,000 resolution with an AGC of 1e5 and a NCE of 35 at a maximum injection time of 250 ms.

### MS data analysis

MS raw data were analyzed using Proteome Discoverer 2.3 (PD 2.3.0.523, Thermo) and the search was performed using the search engine MS Amanda (Dorfer et al., 2014) against a database of the *M. smegmatis* 2019 Uniprot Reference Proteome with contaminants and the BRDT_BD1_ protein added. The database search allowed tryptic peptides with two missed cleavages at a precursor mass tolerance of 5 ppm and 0.02 Da MS2 tolerance. Static alkylation of cysteine and variable oxidation of methionine and TMTpro adducts on lysine and peptide N-termini were considered. Peptides were scored and filtered using Percolator (Kall et al., 2007) to obtain peptides at a 1% false discovery rate. Reporter ions were quantified using the IMP Hyperplex (https://ms.imp.ac.at/?goto=pd-nodes) at a reporter mass tolerance of 10 ppm with a MS2 precursor threshold of 10. The search was performed for each replicate separately over the 10 fractions.

Statistical analysis and data normalization were performed in R. The samples were first normalized for different sample loading by their total sum within each replicate set and then the three TMT replicates were normalized using the Internal Reference Scaling (IRS) method (Plubell et al., 2017). Median alignment was done afterwards by TMM normalization. For each protein, the fold change of TMT-intensities and the corresponding P value (two-tailed Student’s T-test,) were calculated. Permutation-based FDR calculation was used to assess the q-values. The mass spectrometry proteomics data have been deposited to the ProteomeXchange Consortium via the PRIDE (Perez-Riverol et al., 2019) partner repository with the dataset identifier PXD021505.

### Data availability

Coordinates of the ClpC_NTD_:sCym-1 crystal structure have been deposited at the Protein Data Bank (PDBe) under accession code 7AA4. Cryo-EM maps and atomic coordinates have been deposited in the EMDB and PDB with accession codes EMD-11708 for the ClpC tetramer-of-hexamers; EMD-11707 and PDB 7ABR for the single ClpC hexamer composing the assembly. The raw micrographs were submitted to the EMPIAR database (deposition ID: 847). The mass spectrometry proteomics data have been deposited to the ProteomeXchange Consortium via the PRIDE partner repository with the dataset identifier PXD021505.

