## Supplementary Information for "BacPROTACs mediate targeted protein degradation in bacteria"

### SUPPLEMENTARY FIGURES

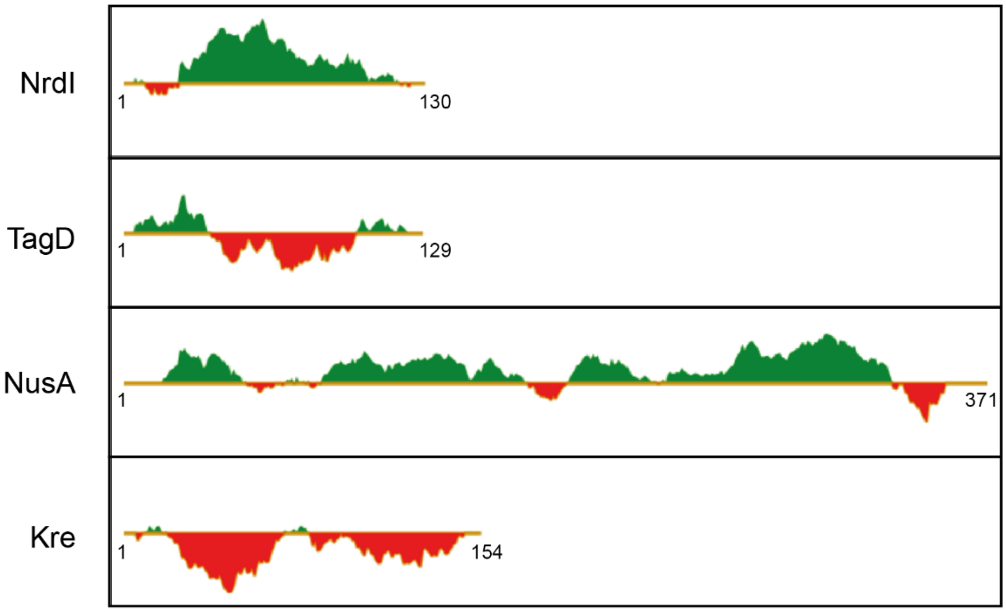


**Figure S1**. **Predicted foldedness of selected model proteins**.

FoldIndex (Prilusky et al., 2005) analysis of *B. subtilis* proteins used in mSA-fusion constructs. Structurally ordered sequences (green) and disordered regions (red) are highlighted. The selected proteins include: NrdI (a flavodoxin-like protein component of ribonucleoside reductases) (Cotruvo and Stubbe, 2010), TagD (glycerol-3-phosphate cytidyltranferase involved in teichoic acid synthesis) (Park et al., 1993), NusA (transcription factor involved in pausing/termination) (Gusarov and Nudler, 2001) and Kre (also known as YkyB, a regulator of the competence transcription factor ComK) (Gamba et al., 2015). PDB structures are available for *B. subtilis* NrdI bound to riboflavin monophosphate (PDB: 1RLJ); *B. subtilis* TagD dimer bound to cytidine-5'-triphosphate (PDB: 1COZ); *Thermotoga maritima* NusA, which has 51% identity to *B. subtilis* NusA (PDB: 1HH2).

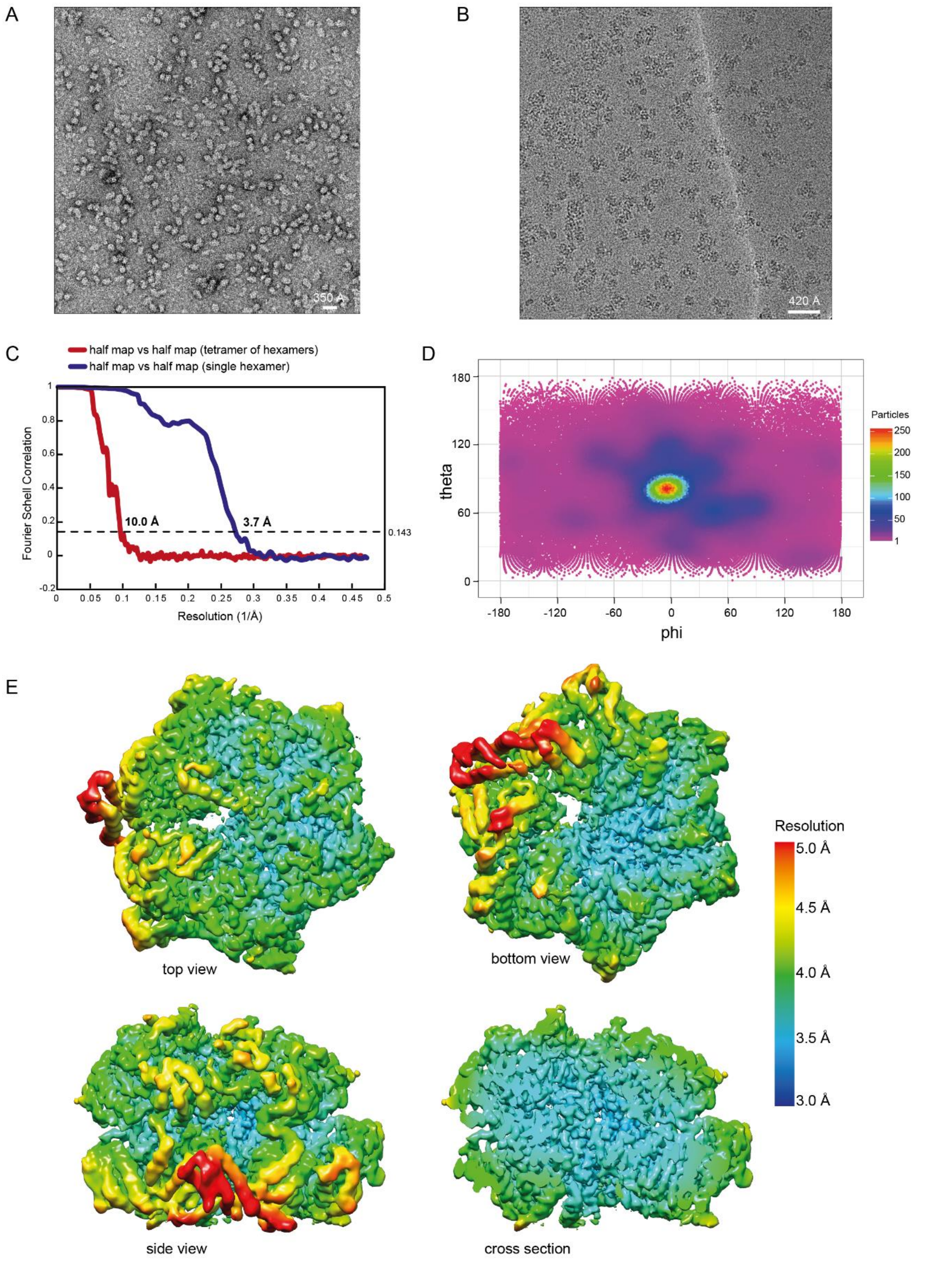


**Figure S2. EM analysis of *B. subtilis* ClpC$_{DWB}$.**

(**A**) Negative stain EM analysis of *B. subtilis* ClpC$_{DWB}$ in the absence of BacPROTAC-1: representative micrograph from the 1,004 collected (scale bar = 350 Å). (**B**) Cryo-EM analysis of *B. subtilis* ClpC$_{DWB}$ in the presence of BacPROTAC-1: a representative micrograph from the 4,455 collected (scale bar = 420 Å). (**C**) FSC curves of the final maps obtained by cryo-EM analysis, showing a resolution of 10 Å for the tetramer of hexamers map and 3.7 Å for the single hexamer map. (**D**) Angular distribution of the particles used to reconstruct the single hexamer map. (**E**) Local resolution map for the single hexamer in different orientations.

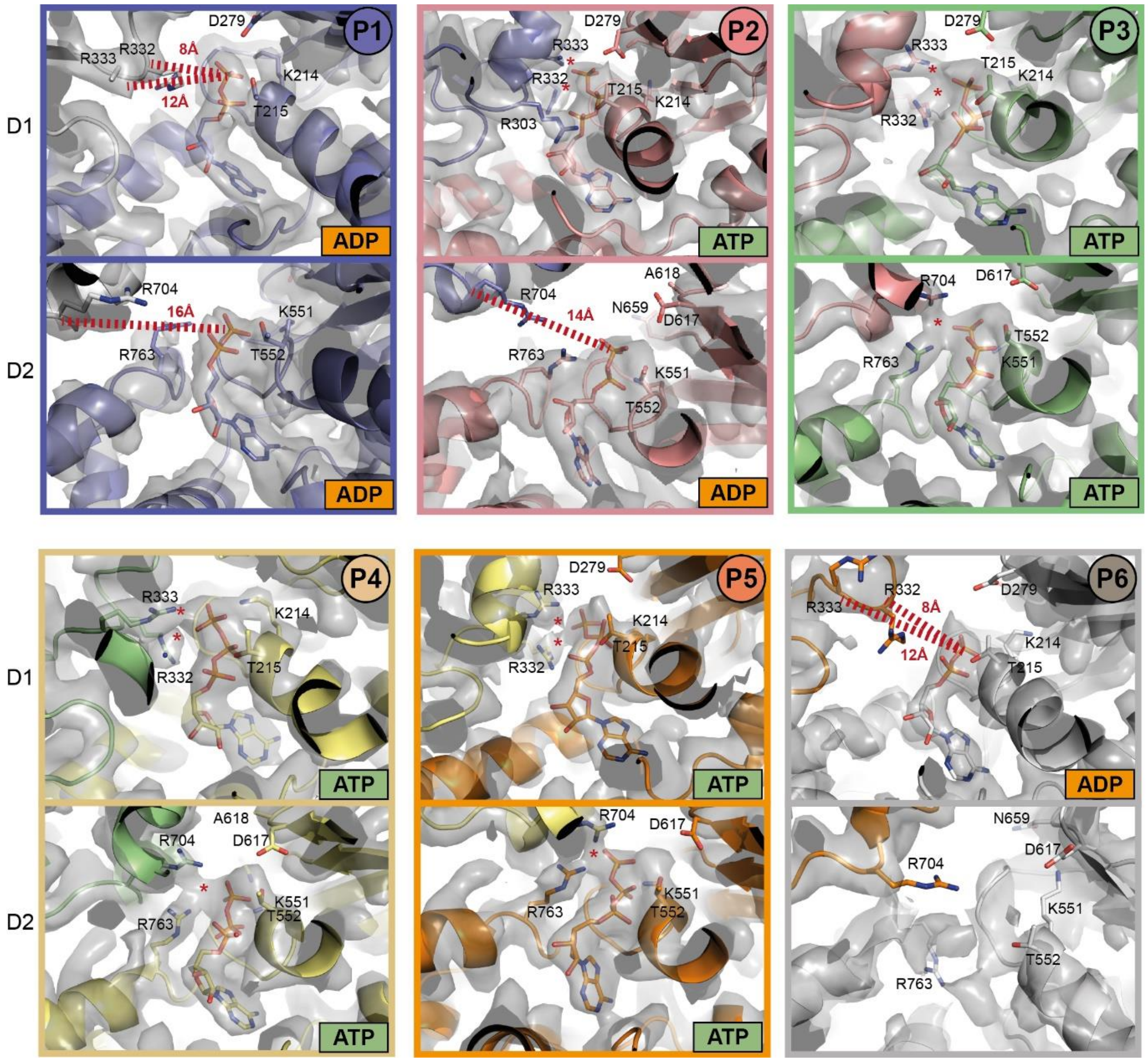


**Figure S3. Nucleotide binding sites of the six ClpC$_{DWB}$ protomers.**

The different panels show the modelled nucleotide in each active site pocket and some of the crucial residues involved in ATP hydrolysis and ATP/ADP interaction. Contacts between ATP γ-phosphate and Arg fingers in D1 (R332-R333) and D2 (R704) are indicated (*) for the ATP-bound sites, while distances between Arg fingers Cα and ADP β-phosphate are shown for ADP-bound sites. The cryo-EM map is represented as grey surface around the modelled protein structure.

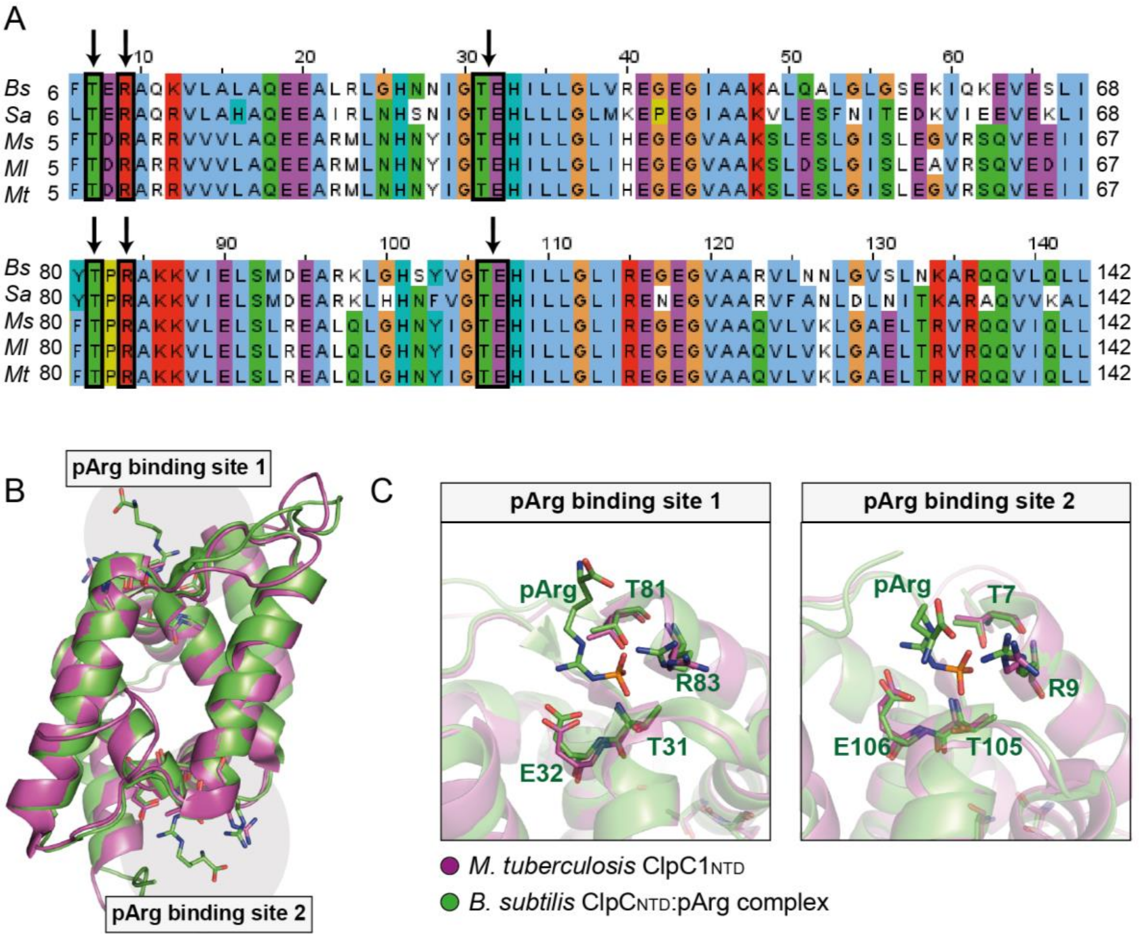


**Figure S4. Conservation of pArg binding sites in Gram-positive bacteria and mycobacteria.**

(**A**) Sequence alignment of ClpC$_{\text{NTDs}}$ from different species (*B. subtilis*, *S. aureus*, *M. smegmatis*, *M. leprae*, *M. tuberculosis*). Residues interacting with pArg are circled in black and marked by an arrow. (**B**) Structure of pArg-bound *B. subtilis* ClpC$_{\text{NTD}}$ (colored green, PDB: 5HBN) superposed with *M. tuberculosis* ClpC1$_{\text{NTD}}$ (colored magenta, PDB: 3WDB). (**C**) Zoomed view of the two pArg binding sites shows that crucial residues interacting with the phospho-guanidino group are conserved in *M. tuberculosis*.

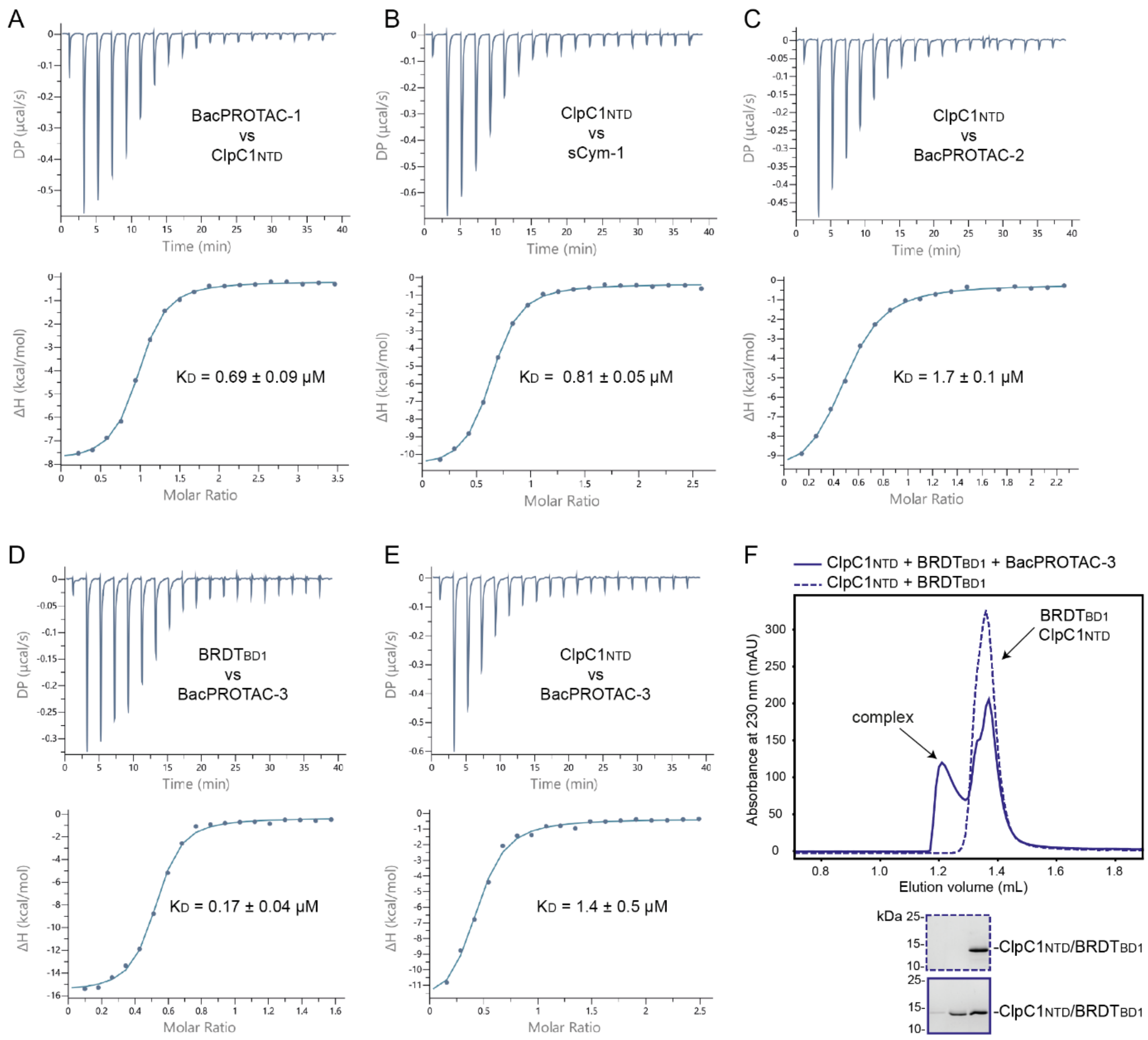


**Figure S5. Characterization of compounds and BacPROTACs binding ClpC1$_{NTD}$.**

(**A**) Representative ITC titration of BacPROTAC-1 (400 µM loaded in the syringe) against ClpC1$_{NTD}$ (22 µM loaded in the cell); reported K$_D$ value represents the average ± standard deviation of three independent measurements. (**B**) Representative ITC titration of ClpC1$_{NTD}$ (406 µM loaded in the syringe) against sCym-1 (30 µM loaded in the cell), reported K$_D$ value represents the average ± standard deviation of three independent measurements. (**C**) Representative ITC titration of ClpC1$_{NTD}$ (356 µM loaded in the syringe) against BacPROTAC-2 (30 µM loaded in the cell), reported K$_D$ value represents the average ± standard deviation of two independent measurements. (**D**) Representative ITC titration of BRDT$_{BD1}$ (124 µM loaded in the syringe) against BacPROTAC-3 (15 µM loaded in the cell); reported K$_D$ value represents the average ± standard deviation of three independent measurements. (**E**) Representative ITC titration of ClpC1$_{NTD}$ (392 µM loaded in the syringe) against BacPROTAC-3 (30 µM loaded in the cell); reported $K_D$ value represents the average ± standard deviation of five independent measurements. (**F**) SEC analysis of a stoichiometric $BRDT_{BD1}$:$ClpC1_{NTD}$ mixture in the presence (solid line) or absence (dashed line) of BacPROTAC-3. The two proteins elute at the same volume also in the absence of BacPROTAC because of their similar size, however BacPROTAC-3 addition mediates formation of an additional peak compatible with the elution volume expected for the $BRDT_{BD1}$:BacPROTAC-3:$ClpC1_{NTD}$ ternary complex. Coomassie stained SDS-PAGE gel of the collected peak fractions is shown. $BRDT_{BD1}$ and $ClpC1_{NTD}$ have identical electrophoretic mobility and are thus not distinguishable on the Coomassie stained gel.

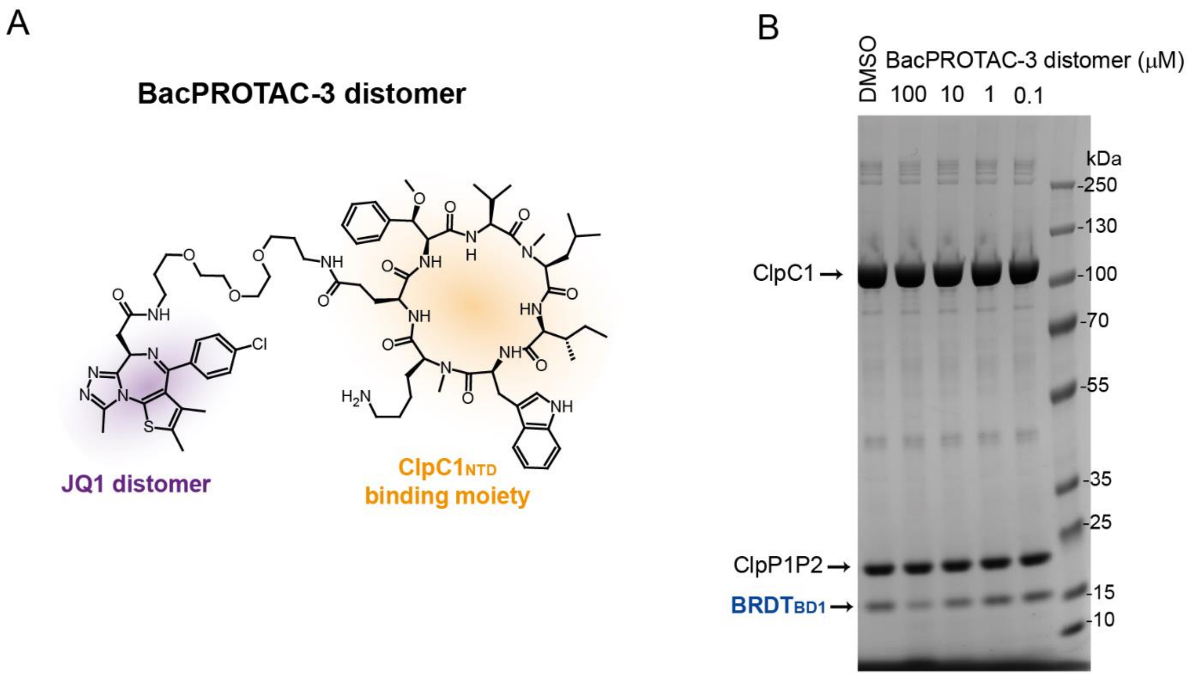


**Figure S6. BacPROTAC-3-induced degradation is stereospecific**.

JQ1 binding to $BRDT_{BD1}$ is stereo-specific (Filippakopoulos et al., 2010). Only the JQ1-(*S)* enantiomer is active (eutomer), while the JQ1-(*R)* enantiomer binds with an approximately 60-fold lower affinity (distomer). (**A**) Chemical structure of the BacPROTAC-3 distomer, synthesized using JQ1-(*R).* (**B**) *In vitro* degradation assay using *M. smegmatis* ClpC1P1P2 analyzed after 2 hours incubation showing that $BRDT_{BD1}$ not significantly degraded in presence of 100-0.1 μM BacPROTAC-3 distomer, in contrast to the BacPROTAC-3 eutomer (**Figure 5B**).

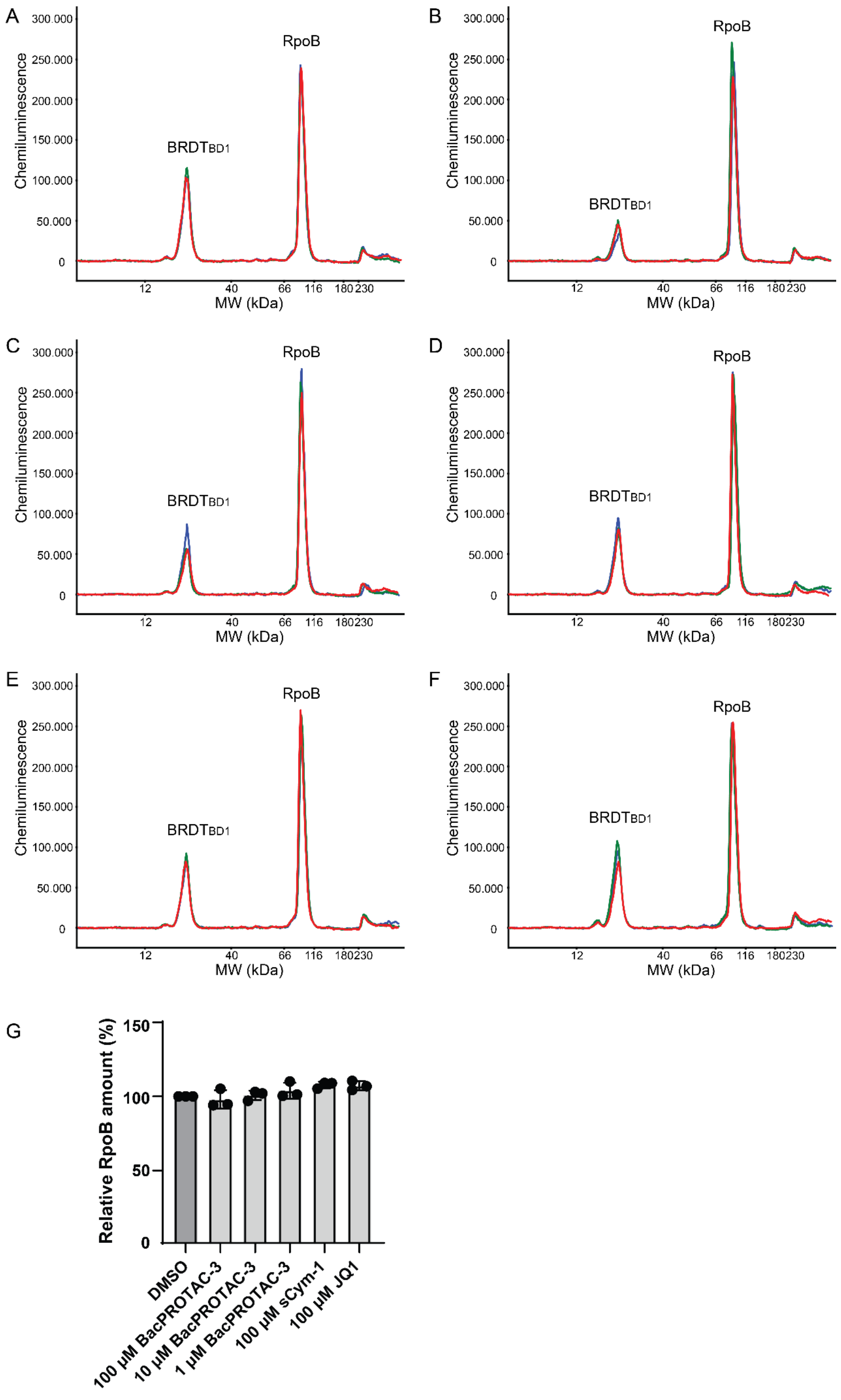


**Figure S7. BRDT$_{BD1}$ levels detected by capillary Western blot upon BacPROTAC treatment of *M. smegmatis* cells.**

Electropherograms showing intensity of the chemiluminescent signal plotted against the apparent molecular weight detected using anti-BRDT and anti-RpoB antibodies.

(**A-F**) Detected peaks for lysates of a *M. smegmatis* culture expressing $BRDT_{BD1}$ after each treatment (30 minutes) in triplicate: (**A**) DMSO, (**B**) 100 µM BacPROTAC-3, (**C**) 10 µM BacPROTAC-3, (**D**) 1 µM BacPROTAC-3, (**E**) 100 µM sCym-1, (**F**) 100 µM JQ1. (**G**) Bar chart showing quantification of detected RpoB peaks (loading control) from three independent experiments normalized to DMSO treatment (dark grey bar) and plotted as mean ± standard deviation. Quantification of the $BRDT_{BD1}$ peak is shown in **Figure 5D**.

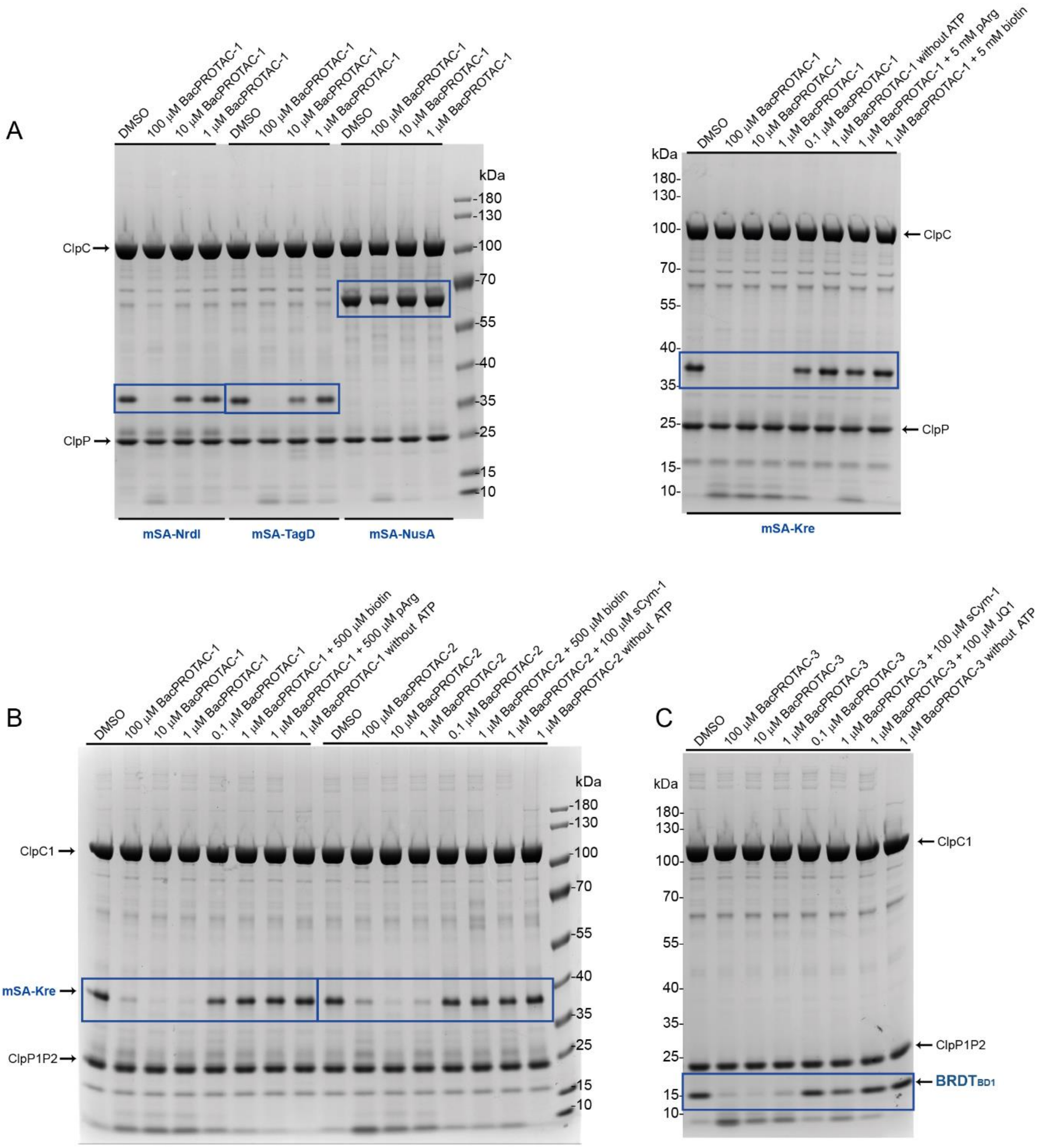


**Figure S8. Uncropped assay gels.** Uncropped Coomassie stained SDS-PAGE gels shown in (**A**) **Figure 1G** and **1H**; (**B**) **Figure 4B** and **4H**; (**C**) **Figure 5B**.

### SUPPLEMENTARY TABLES

**Table S1: Crystallographic analysis of ClpC1$_{NTD}$:sCym-1 complex. Data collection and refinement statistics**

| | |
|---|---|
| PDB ID | 7AA4 |
| Space group | *P*1 |
| Cell dimensions | |
| *a*, *b*, *c* (Å) | 31.35, 33.68, 35.81 |
| *α*, *β*, *γ* (°) | 86.178, 94.216, 103.176 |
| Resolution (Å)[a,b] | 25 – 1.68 (1.72 – 1.68) |
| $R_{meas(I)}$ | 0.062 (0.112) |
| *I/σ* (*I*) | 22.6 (12.5) |
| $CC_{1/2}$ | 0.998 (0.992) |
| Completeness (%) | 93.8 (86.5) |
| Redundancy | 5.8 (5.4) |
| Resolution (Å) | 25 – 1.68 |
| No. reflections | 15,165 |
| $R_{work}$ / $R_{free}$ | 17.0 / 20.3 |
| No. atoms | |
| protein | 1254 |
| ligand | 66 |
| water | 192 |
| *B* factors | |
| protein | 11.9 |
| ligand | 13.5 |
| water | 23.0 |

| | |
|---|---|
| R.m.s. deviations | |
| Bond lengths (Å) | 0.006 |
| Bond angles (°) | 0.789 |

[a]Values in parentheses are for highest-resolution shell.

[b]Due to experimental constraints the resolution needed to be truncated to this resolution.

**Table S2. Amino acid sequences of mSA fusion proteins.**

| **Construct name** | **Amino acid sequence** |
|---|---|
| mSA | MHHHHHHSSGVDLGTENLYFQSSQDLASAEAGITGTWYNQSGSTFTVTAGADGNLTGQYENRAQGTGCQNSPYTLTGRYNGTKLEWRVEWNNSTENCHSRTEWRGQYQGGAEARINTQWNLTYEGGSGPATEQGQDTFTKVKPSAASGSGSGSGSGS |
| mSA-Kre | MHHHHHHSSGVDLGTENLYFQSSQDLASAEAGITGTWYNQSGSTFTVTAGADGNLTGQYENRAQGTGCQNSPYTLTGRYNGTKLEWRVEWNNSTENCHSRTEWRGQYQGGAEARINTQWNLTYEGGSGPATEQGQDTFTKVKPSAASGSGSGSGSMDDHAYTKDLQPTVENLSKAVYTVNRHAKTAPNPKYLYLLKKRALQKLVKEGKGKKIGLHFSKNPRFSQQQSDVLISIGDYYFHMPPTKEDFEHLPHLGTLNQSYRNPKAQMSLTKAKHLLQEYVGMKEKPLVPNRQQPAYHKPVFKKLGESYF |
| mSA-Kre (Cryo-EM structure determination) | MSQDLASAEAGITGTWYNQSGSTFTVTAGADGNLTGQYENRAQGTGCQNSPYTLTGRYNGTKLEWRVEWNNSTENCHSRTEWRGQYQGGAEARINTQWNLTYEGGSGPATEQGQDTFTKVKPSAASGSGSGSGSGSGSGSGSGSDDHAYTKDLQPTVENLSKAVYTVNRHAKTAPNPKYLYLLKKRALQKLVKEGKGKKIGLHFSKNPRFSQQQSDVLISIGDYYFHMPPTKEDFEHLPHLGTLNQSYRNPKAQMSLTKAKHLLQEYVGMKEKPLVPNRQQPAYHKPVFK KLGESYFHHHHHH |
| mSA-NrdI | MHHHHHHSSGVDLGTENLYFQSSQDLASAEAGITGTWYNQSGSTFTVTAGADGNLTGQYENRAQGTGCQNSPYTLTGRYNGTKLEWRVEWNNSTENCHSRTEWRGQYQGGAEARINTQWNLTYEGGSGPATEQGQDTFTKVKPSAASGSGSGSGSGSVVQIIFDSKTGNVQRFVNKTGFQQIRKVDEMDHVDTPFVLVTYTTNFGQVPASTQSFLEKYAHLLLGVAASGNKVWGDNFAKSADTISRQYQVPILHKFELSGTSKDVELFTQEVERVVTKSSAKMDPVK |
| mSA-TagD | MHHHHHHSSGVDLGTENLYFQSSQDLASAEAGITGTWYNQSGSTFTVTAGADGNLTGQYENRAQGTGCQNSPYTLTGRYNGTKLEWRVEWNNSTENCHSRTEWRGQYQGGAEARINTQWNLTYEGGSGPATEQGQDTFTKVKPSAASGSGSGSGSGSMKKVITYGTFDLLHWGHIKLLERAKQLGDYLVVAISTDEFNLQKQKKAYHSYEHRKLILETIRYVDEVIPEKNWEQKKQDIIDHNIDVFVMGDDWEGKFDFLKDQCEVVYLPRTEGISTTKIKEEIAGL |
| mSA-NusA | MHHHHHHSSGVDLGTENLYFQSSQDLASAEAGITGTWYNQSGSTFTVTAGADGNLTGQYENRAQGTGCQNSPYTLTGRYNGTKLEWRVEWNNSTENCHSRTEWRGQYQGGAEARINTQWNLTYEGGSGPATEQGQDTFTKVKPSAASGSGSGSGSGSMSSELLDALTILEKEKGISKEIIIEAIEAALISAYKRNFNQAQNVRVDLNRETGSIRVFARKDVVDEVYDQRLEISIEEAQGIHPEYMVGDVVEIEVTPKDFGRIAAQTAK |

| | |
|---|---|
| | QVVTQRVREAERGVIYSEFIDREEDIMTGIVQRLDNKFIYVSLGKIEALLPVNEQM<br>PNESYKPHDRIKVYITKVEKTTKGPQIYVSRTHPGLLKRLFEIEVPEIYDGTVELKS<br>VAREAGDRSKISVRTDDPDVDPVGSCVGPKGQRVQAIVNELKGEKIDIVNWSSD<br>PVEFVANALSPSKVLDVIVNEEEKATTVIVPDYQLSLAIGKRGQNARLAAKLTGW<br>KIDIKSETDARELGIYPRELEEDDEPLFTEPETAESDE |

**Movie S1. Cryo-EM structure of activated ClpC in complex with substrate.**

The movie shows the map of the tetramer of ClpC hexamers (10 Å) and the map obtained refining a single ClpC hexamer (3.7 Å). Both maps are colored by ClpC protomer. Continuous density is observed for a substrate peptide (colored yellow) along the ClpC channel, surrounded by Tyr-containing pore loops, as described in the main text.
